# Spatio-temporal organization of network activity patterns in the hippocampus

**DOI:** 10.1101/2023.10.17.562689

**Authors:** Vítor Lopes-dos-Santos, Demi Brizee, David Dupret

**Affiliations:** Medical Research Council Brain Network Dynamics Unit, Nuffield Department of Clinical Neurosciences, University of Oxford, Oxford, UK

## Abstract

The hippocampus is a layered brain network, composed of diverse cell types arranged in multiple microcircuits with anatomically structured inputs, all working in concert to support memory. This intricate organization generates a myriad of electrophysiological signatures. While specific aspects of these activity patterns have been explored, a comprehensive understanding of the hippocampal layer-embedded dynamics remains elusive. Here, we developed a low-dimensional manifold to capture electrophysiological patterns, mapping their anatomical trajectory along the CA1-to-DG axis, and distinguishing layers based on sharp-wave and theta profiles. This profiling led to the characterization of selective theta-nested gamma signatures for each layer. It further revealed spike patterns associated with gamma rhythms, which highlight specific firing motifs between principal cells and interneurons, and differential firing properties within pyramidal sub-layers. These findings support a holistic understanding of the spatio-temporal activity patterns across hippocampal layers for unraveling the network operations that drive memory-guided behavior.

## Introduction

The hippocampus is a neural network at the nexus of the brain circuitry dedicated to processing mnemonic information (Andersen, 2007; Eichenbaum, 2000; Van Strien et al., 2009). The culmination of the vast computations within this network is embodied in the diverse firing patterns of the CA1 pyramidal cell ensemble (Buzsáki, 2010; O’Keefe and Nadel, 1978). This output, shaped by numerous inputs from both inside and outside the hippocampus, is further refined by local processes to guide memory-related behavior (Klausberger and Somogyi, 2008; Mizuseki et al., 2009). These neural inputs exhibit an orderly arrangement along the somato-dendritic axis of CA1 principal cells (Spruston, 2008). The structured radial organization across the distinct anatomical layers of the hippocampus results in multiple electrophysiological patterns, observable in the local field potentials (LFPs) recorded along that axis. Reaching a comprehensive understanding of these network activity signatures would shed light on the intricately woven spatio-temporal processes that dynamically support the pivotal contribution of the hippocampus to memory and behavior.

The hippocampal network patterns are behavioral state-dependent and grounded in cellular processes. During sleep and immobility periods, prominent phenomena include the sharp-wave ripple (SWR) complex in the *Cornu Ammonis* (CA) and dentate spikes in the dentate gyrus (DG). Dentate spikes represent intermittent, large amplitude events that are recorded in the LFPs of the DG granule cell layer and associated with entorhinal cortex inputs (Bragin et al., 1995b; Penttonen et al., 1997). SWRs arise from a current sink in the CA1 radiatum layer (the sharp-wave) caused by CA3 inputs, which culminates in brief high-frequency (100 – 250 Hz) oscillations (ripples) within the CA1 pyramidal layer (Buzsáki, 1986; Csicsvari et al., 1999; Ylinen et al., 1995). Deciphering the neural mechanisms behind these phenomena has generated a wealth of research, notably elucidating the role of SWRs in offline memory stabilization and awake learning processes (Buzsáki, 2015; Ego-Stengel and Wilson, 2009; Girardeau et al., 2009; van de Ven et al., 2016). During active exploration periods, the hippocampus predominantly exhibits theta (5 – 12 Hz) oscillations (Buzsáki, 2005; Vanderwolf, 1969). This rhythm not only coordinates multiple hippocampal circuits but also orchestrates their interaction with extrahippocampal regions (Benchenane et al., 2010; Hyman et al., 2005; Jones and Wilson, 2005; Siapas et al., 2005; Sirota et al., 2008). A prominent feature of the theta rhythm is its nested gamma oscillations (25 – 150 Hz), a set of diverse, rapid oscillatory bursts that reflect the activity of distinct neural circuits and pathways across the hippocampal formation (Bragin et al., 1995a; Colgin et al., 2009; Fernandez-Ruiz et al., 2023; Tort et al., 2009). Together, these distinct network patterns provide a nuanced lens into the versatile functional dynamics of the hippocampus across varying behavioral states.

In this study, to explore the temporal architecture of hippocampal layer-related network activity, we developed a low-dimensional feature manifold embedding electrophysiological patterns recorded in freely behaving mice. We demonstrate that individual LFP signals along the CA1-to-DG axis carry sufficient information within their theta and sharp-wave profiles to discern the radially organized hippocampal layers. Furthermore, we observe that each discerned layer displays a distinct theta-nested gamma profile, underscoring the coexistence of multiple gamma oscillations within the hippocampus. We analyze the relationships between these gamma oscillations and the firing patterns of CA1 principal cells and interneurons, uncovering distinct firing motifs associated with gamma rhythms. Leveraging our LFP-embedded manifold, we further categorize CA1 principal cells into deep and superficial pyramidal sub-layer subpopulations, showing differences in their firing properties. Altogether, these results reveal the complex temporal architecture of hippocampal layer-related activity dynamics.

## Results

### Identifying individual hippocampal layers along the CA1-to-DG axis

We first tested whether electrophysiological patterns of the LFPs could distinguish between the radially organized anatomical layers from CA1 to DG in the dorsal hippocampus. For this, we implanted linear silicon probes in six mice along the somato-dendritic axis of CA1 principal cells, extending up to the first DG blade (**Figure 1A**). This configuration provided simultaneous LFP recordings across the hippocampal layers as mice explored open fields and underwent sleep/rest periods. From these recordings, we examined network activity patterns within each recording channel and their cross-channel interactions, including CSD analyses. We used three electrophysiological patterns (**Figure 1B**): theta oscillations (5 – 12 Hz) prevalent during exploratory behavior, as well as sharp-wave ripples (SWRs; 100 – 250 Hz) and dentate spikes (DS1 and DS2), both observed during sleep/rest. We determined the CA1 pyramidal cell layer channel as the one with the highest ripple power (**Figure 1C**). The radiatum layer channel was identified as the one exhibiting the strongest current sink coinciding with the peak of ripple amplitude in the CA1 pyramidal layer (Buzsáki et al., 1983; Karalis and Sirota, 2022; Sullivan et al., 2011; Ylinen et al., 1995). The hippocampal fissure channel was marked by the highest theta power (Brankack et al., 1993; Buzsáki et al., 1986; Gerbrandt and Fowler, 1980; Green and Rawlins, 1979), while the lacunosum-moleculare channel was identified as being 40–50 μm above the fissure. The outer and mid-molecular layers of the DG corresponded to channels with the deepest current sinks for DS1 and DS2 events, respectively (Bragin et al., 1995b; Karalis and Sirota, 2022). We recognized the inner molecular layer by the secondary current sink associated with the radiatum sharp-wave event (Buzsáki, 1986; Karalis and Sirota, 2022). The DG granular layer channel was determined from the strongest current source of DS beneath the molecular layer channels (Bragin et al., 1995b; Karalis and Sirota, 2022; Penttonen et al., 1997; Senzai and Buzsáki, 2017). Thus, we used electrophysiological patterns to map the distinct hippocampal layers along the radial CA1-to-DG axis. These tempo-spatial patterns were consistent across mice and silicon probe layouts (**Supplementary** Figure 1).

**Figure 1:**
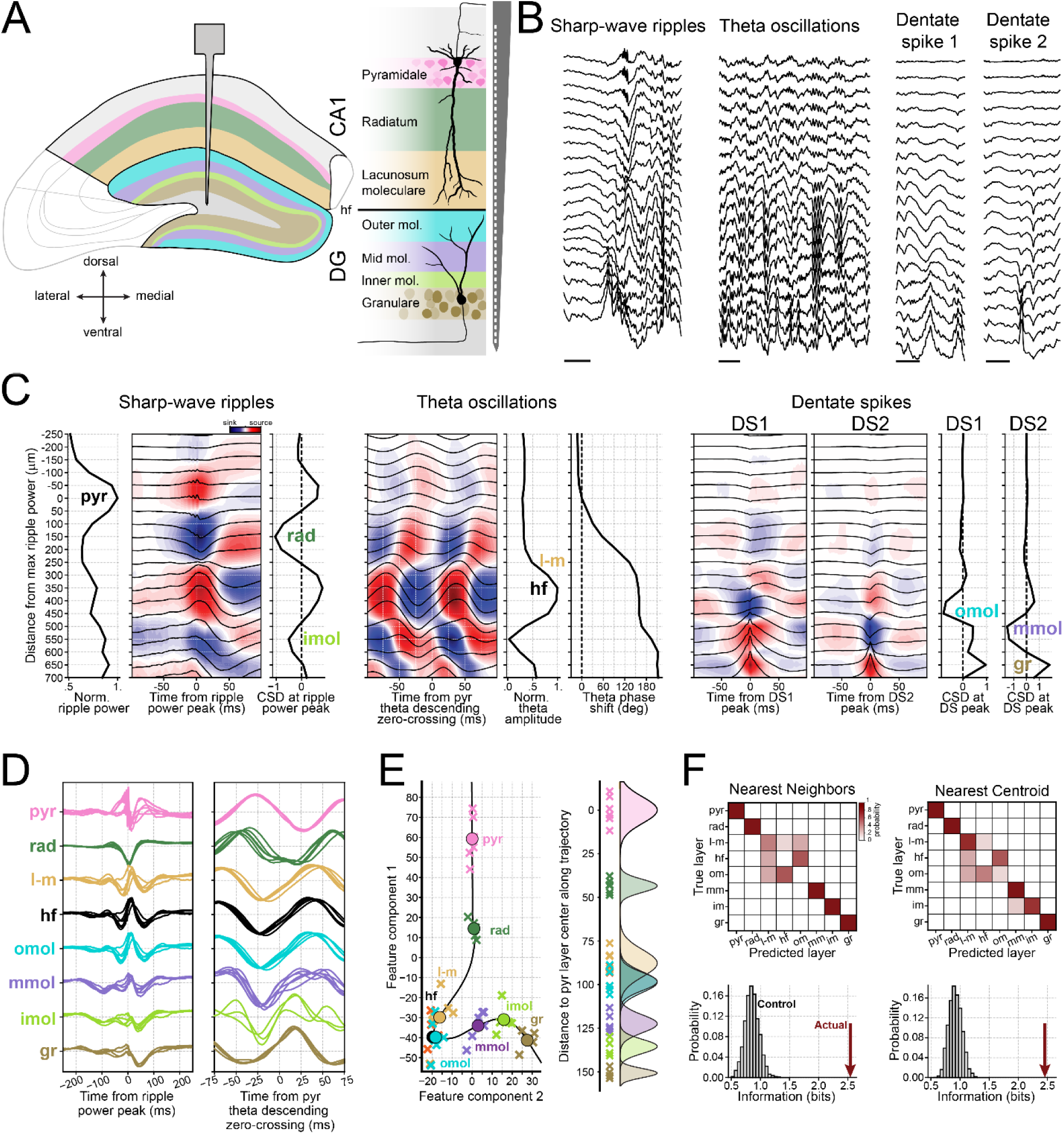
Identification of hippocampal layers using electrophysiological patterns. **(A)** Schematic of the hippocampal layers simultaneously recorded using a silicon probe implanted along the CA1-DG axis. **(B)** Sample raw data showing the three network events used for layer identification. Scale bars: 100 ms. **(C)** Example activity profiles for layer determination from a representative mouse (Mouse 1; See Supplementary Figure 1 for the other mice). From left to right: The sharp-wave ripple panels display ripple power across layers, CSD analysis of LFPs aligned to ripple power peaks, and CSD values at the ripple power peak. The theta oscillation panels display the CSD for LFPs aligned to the pyramidal layer’s descending zero-crossing, alongside normalized theta amplitude and phase shift across layers. The phase shifts are relative to the phase of the pyramidal layer. The dentate spike panels depict CSD analysis for DS1 and DS2 with their respective CSD values at their peak. **(D)** Sharp-wave and theta waveforms for each manifold-defined layer for all mice (each waveform represents one mouse). **(E)** Left: LFP-based feature manifold with individual layers denoted by crosses for each mouse, color-coded as in (D), with circles representing layer averages. Right: Linear trajectory of the manifold, with individual projections determined by their closest trajectory point. See also Supplementary Figure 2. **(F)** Top: Confusion matrices for a Nearest Neighbor Classifier and a Nearest Centroid Classifier used in determining the layer of specific feature manifold data points. For testing any given point, all other points were used in training sets. Entries in the matrices represent the likelihood of predicting a particular layer given the true layer. Bottom: mutual information between the actual layers and the predicted layers by each classifier (red arrows) along with their control distribution computed by shuffling layers in the training sets. Layers: *pyr*, pyramidale; *rad*, radiatum; *l-m*, lacunosum-moleculare; *hf*, hippocampal fissure; *omol*, outer molecular; *mmol*, mid molecular; *imol*, inner molecular; *gr*, granular.

The layer assignment above benefited from linear silicon probes that enabled simultaneous recordings from uniformly spaced channels along the CA1-DG axis. Next, we sought to determine how much information for layer assignment could be drawn only from the theta phase reversal and changes in the sharp-wave waveform on a per-channel basis. For this, we employed ISOMAP embedding to represent data from both theta and sharp-wave waveforms across hippocampal layers and mice (**Figure 1C,D** and **Supplementary** Figure 2) into a 2-dimensional feature space (**Figure 1E**). This dimensionality reduction method revealed that channels recording from the same hippocampal layer tend to form separate clusters from those of other layers. Additionally, the derived feature trajectory naturally connected anatomically adjacent layers, even without explicitly feeding this layer adjacency information into the manifold embedding.

To further assess the electrophysiological distinguishability of hippocampal layers, we employed nearest neighbor and nearest centroid-based classifiers, training them to recognize each individual layer from its feature components, using a leave-one-out cross-validation (**Figure 1F**). For the pyramidal, radiatum, mid molecular, and granular layers, this approach achieved optimal classification with both classifiers. The inner molecular layer achieved near-optimal classification. While the lacunosum-moleculare layer, hippocampal fissure, and outer molecular layer could be effectively differentiated from all other layers, splitting this group of three coalescing layers was not possible (**Figure 1F**; top panels). To quantify how much layer-specific information can be obtained from the feature manifold, we measured the Shannon information between actual and predicted layers by the classifiers. These values significantly exceeded those from control sets with shuffled labels (**Figure 1F**; bottom panels). Together, these findings demonstrate that LFP-based features can discern different anatomical layers spanning the hippocampal CA1-DG axis.

### Laminar profile of CA1-DG theta-nested gamma oscillations

Theta oscillations influence broad brain networks, while hippocampal gamma oscillations are believed to relate to finer-grained neural pathways (Colgin et al., 2009; Fernandez-Ruiz et al., 2023). Thus, analyzing gamma patterns across layers could aid reaching a comprehensive understanding of the intricate local dynamics and functional microarchitecture of the hippocampus network. Accordingly, we generated theta-gamma profiles for each layer identified in our LFP manifold embedding (**Figure 1E**), using two evaluations (**Figure 2A** and **Supplementary** Figure 3A). First, we gauged the amplitude of individual frequencies within the gamma range using the theta rhythm from the CA1 pyramidal layer as a reference (**Figure 2A**; left panel for each layer). This allowed comparing amplitudes of gamma frequencies within each layer. Second, we normalized the amplitude of each frequency by its minimum value (**Figure 2A**; right panel for each layer). This rendered a perspective on the amplitude modulation of various gamma oscillations in relation to the theta phase, regardless of their individual magnitudes.

**Figure 2:**
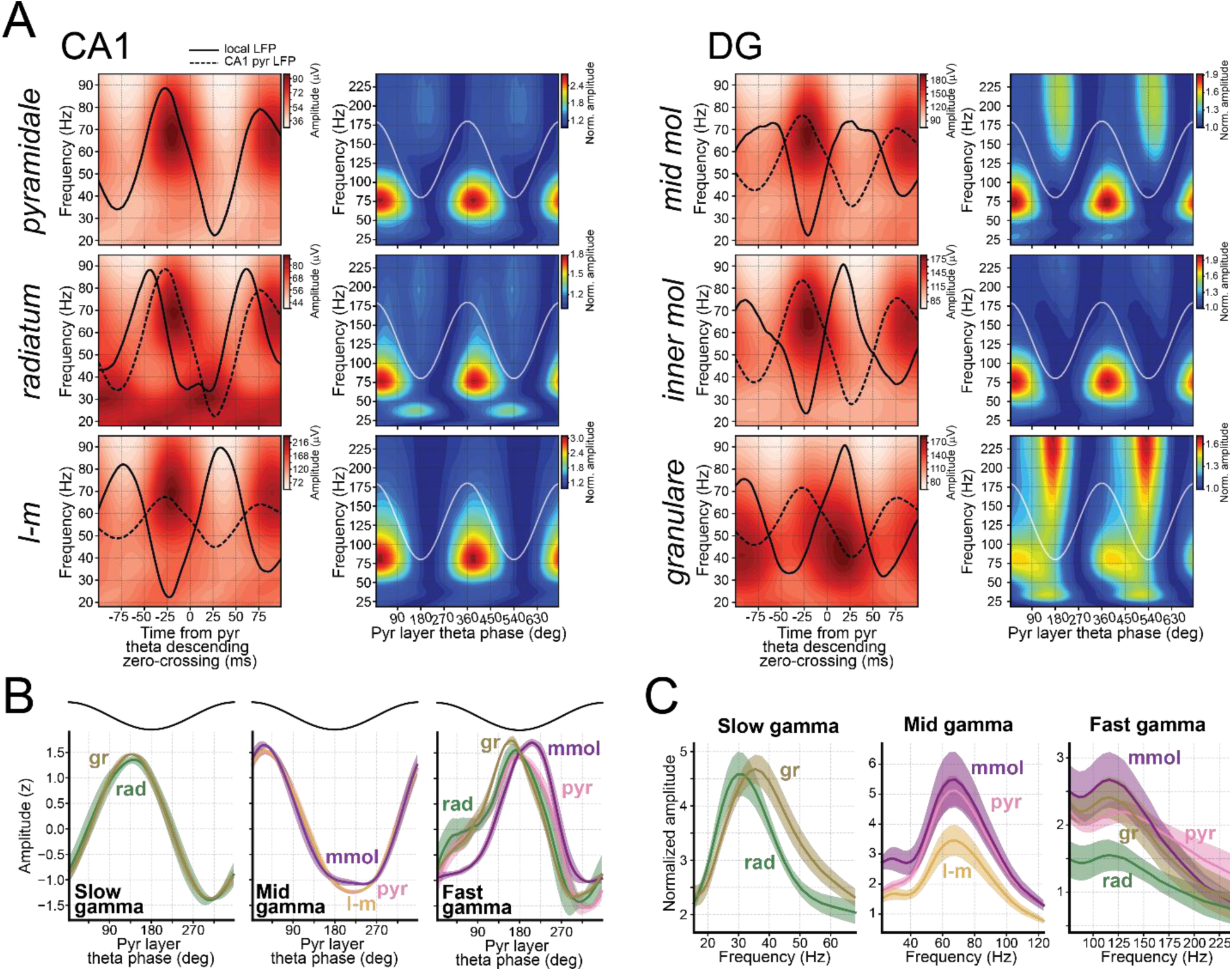
Theta-nested gamma profiles across individual CA1-to-DG layers. **(A)** Each layer features two associated panels displaying theta-nested gamma amplitudes. In the left panels, both local (solid lines) and CA1 pyramidal layer LFPs (dashed lines) are shown, aligned to the descending zero-crossing of the pyramidal layer theta reference. These are juxtaposed with amplitude of gamma frequencies aligned similarly. The right panels depict the amplitude of these gamma frequencies, normalized with respect to the pyramidal layer theta phase. Amplitude normalization was done by dividing by the minimum amplitude for each frequency. The reference cosine is overlayed in white. The layers shown here are from one example mouse (see Supplementary Figure 3A for the other mice). **(B)** Z-scored amplitude for designated gamma bands within the assorted layers in relation to the pyramidal layer theta phase. A cosine atop provides a phase reference. **(C)** Normalized power spectra (see *Determination of gamma main frequencies* in the Method section for details) calculated for each gamma band in specific layers. In both B and C, solid lines denote mean across mice while shaded regions represent 95% confidence intervals.

In CA1, we found mid gamma (range: ∼50 – 90 Hz) oscillations across all layers (**Figure 2A**), with the maximum amplitude occurring just after the CA1 pyramidal layer theta peak (**Figure 2A,B**). The pyramidal layer was characterized by a fast gamma oscillation (range: ∼100 – 250 Hz) that increased at the theta trough (**Figure 2A,B**). The radiatum layer featured a slow gamma component at the descending phase of theta, near the trough (range: ∼25 – 45 Hz; **Figure 2A,B**), along with a slightly slower fast gamma than that in the pyramidal layer (**Figure 2A,B**). The lacunosum-moleculare layer showed mid gamma oscillations, without clear slow or fast gamma components (**Figure 2A**). These findings were reproducible across mice with different silicon probe implants (**Supplementary** Figure 3A).

In the DG, two distinct fast gamma components emerged. The first was evident in the mid-molecular layer, emerging just after the CA1 pyramidal theta trough; while the second was localized to the granular layer, just prior to the CA1 pyramidal theta trough (**Figure 2A,B** and **Supplementary** Figure 3A). The granular layer also displayed a slow gamma component that was aligned with the slow gamma observed in the radiatum layer (**Figure 2A,B** and **Supplementary** Figure 3A). Like CA1, all DG layers displayed mid gamma oscillations.

Additionally, a beta component (range: ∼18 – 35 Hz) was observed in mid-molecular layer channels, peaking post the CA1 pyramidal theta peak, coinciding in theta phase with mid gamma oscillations (**Supplementary** Figure 3A). This beta component occupies the lower gamma frequencies but was distinct from the slow gamma oscillations. While the slow gamma oscillations partially overlap with the beta range, they distinctly emerge in the radiatum, with their amplitude achieving a maximum near the pyramidal theta trough.

Assigning a precise frequency range for gamma oscillations is challenging. This issue is heightened for fast gamma oscillations, which exhibit significantly lower amplitudes than mid gamma oscillations and can therefore be overshadowed in non-normalized spectrograms. On the other hand, normalized spectrograms may distort the frequency range of overlapping gamma components (**Supplementary** Figure 3B). To circumvent this, we identified bursts of filtered LFP signals within their corresponding bands and calculated the average broadband LFP spectrograms centered around the maximum amplitude of these bursts (see Methods). This indicated a frequency peak of ∼31 Hz for slow gamma in the radiatum, and ∼35 Hz in the granular layer (**Figure 2C**; left panel). Mid gamma displayed a main frequency of ∼68 Hz across all layers (**Figure 2C**; mid panel). Fast gamma oscillations exhibited a peak frequency of ∼130 Hz in the pyramidal layer, and ∼120 Hz in the radiatum, molecular layer, and granular layer (**Figure 2C**; right panel).

In sum, these observations obtained from the signals recorded using linear silicon probes consistently revealed slow gamma oscillations in the radiatum and granular layers, as well as a widespread mid-gamma component across all CA1-to-DG layers and mice. Notably, this layer profiling also suggested the existence of four fast gamma components across the pyramidal, radiatum, mid molecular, and granular layers. Finally, the consistency observed in the gamma profiles across mice provides important cross-validation to the hippocampal layer delineation, as gamma oscillations were not utilized in the LFP feature manifold.

### Laminar signatures retrieved in independently moveable tetrode recordings

To assess the generalizability of the obtained hippocampal activity layer profiling across mice and recording techniques, we employed the manifold analysis to guide tetrode placement in additional mice implanted with independently moveable tetrodes. For each recording, we lowered tetrodes individually in a stepwise manner, acquiring their signals at each step while maintaining one tetrode in the pyramidal layer as a reference channel for ripple power peak and theta phase. Subsequent histological evaluation validated the accuracy of the manifold-derived predictions. Notably, tetrodes identified in the radiatum, lacunosum-moleculare, mid molecular, and granular regions indeed corresponded to their respective anatomical layers (**Figure 3**). Further, we found that these (manifold-positioned and anatomically confirmed) tetrodes displayed gamma signatures consistent with those obtained in the corresponding layers from silicon probe recordings (**Figure 3** and **Supplementary** Figure 4). Specifically, fast gamma oscillations were consistently observed in the pyramidal, radiatum, and mid molecular layers, occurring just after the pyramidal theta trough. In the granular layer, they preceded the pyramidal theta trough. Importantly, the radiatum fast gamma was more prominent below the “center” of this layer manifold coordinate, that is in a distal part of this layer (**Supplementary** Figure 5). Slow gamma components were also consistently present in the radiatum and granular layer channels with tetrodes, as they were with silicon probes. A beta component was also primarily found in the molecular layer channels. Mid gamma oscillations were also observed across all layers with tetrodes, as with silicon probes. These findings support the presence of distinct theta-gamma profiles locally in each of hippocampal layers individually recorded in the tetrode dataset, in line with the laminar recordings of the silicon probe dataset.

**Figure 3:**
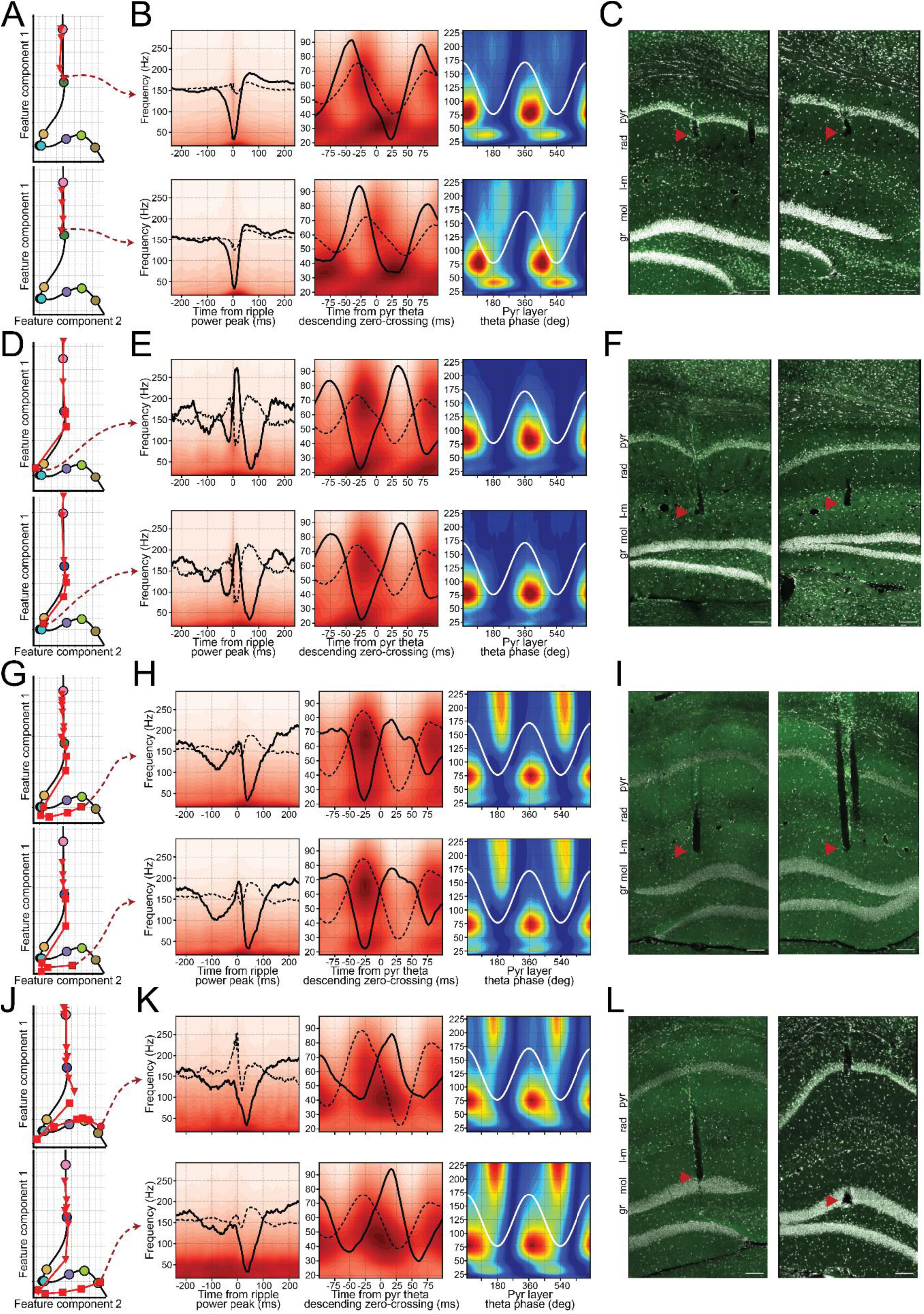
Validation of LFP profiles using tetrode placement in the feature manifold. (**A-C**) Individual tetrode placement in the radiatum layer, guided by electrophysiological patterns. (A) Corresponding trajectories in the feature manifold where each triangle represents a tetrode’s projection for a recording session (with connecting lines indicating the session sequence). Each panel is an example tetrode. (B) shows the sharp-wave and theta-gamma profiles of these tetrodes immediately before perfusion. Left: SWR waveform of the channel (solid) versus reference pyramidal layer (dashed); heatmap displays amplitude across frequencies. Mid and right panels are theta-gamma profiles as the ones in Figure 2. (C) shows the histological confirmation of these tetrodes in radiatum guided by manifold projections. Red arrowheads denote the tip of the tetrodes. Scale bars: 100 μm. (**D-F**) Same analysis but for tetrodes targeting the lacunosum-moleculare layer. Each marker signifies same-day sessions, with triangles and squares indicating consecutive days. There was no tetrode adjustment between the final session of one day and the initial session of the next. (**G-I**) Same analysis but for tetrodes targeting the DG molecular layer. (**J-L**) Same analysis but for tetrodes targeting the DG granular layer. Circles indicate a third recording day.

### Differential spiking correlates of layer-defined gamma oscillations

Gamma oscillations are thought to represent excitatory postsynaptic potentials influencing local spiking activity. Investigating the temporal interplay between neuronal spiking and gamma oscillations offers a window into their foundational circuit mechanisms and significance. In line with this, we observed a strong relationship between the phase of gamma oscillations and the timing of CA1 spikes in both principal cells and interneurons (**Figure 4**). Aligning the activity of both cell populations by a single trough of radiatum slow gamma per theta cycle showed clear slow gamma-paced spiking spanning at least 5 cycles in both populations (**Figure 4A,B**). Furthermore, the mean phase for both CA1 principal cells and interneurons was consistent at the radiatum slow gamma troughs (**Figure 4C**). The same analysis applied to mid gamma oscillations recorded in the lacunosum-moleculare layer revealed 3 cycles of mid gamma-paced modulation in the spiking activity of CA1 cells (**Figure 4D,E**). We however noted that while CA1 principal cells consistently fired preferentially around lacunosum-moleculare mid gamma ascending phase, interneurons displayed varied firing phases from the mid gamma trough to its peak (**Figure 4F**).

**Figure 4:**
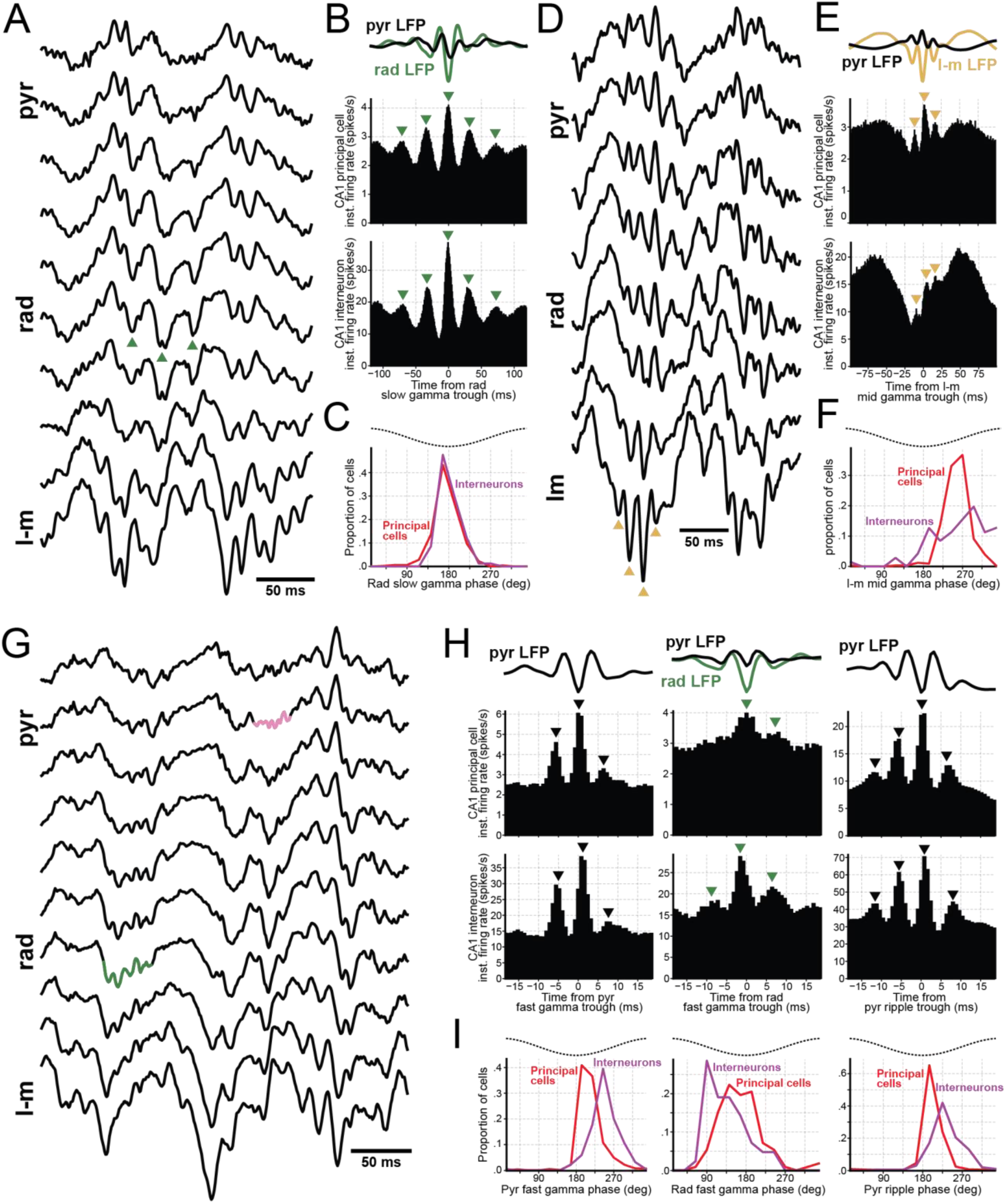
Spiking correlates of CA1 principal cells and interneurons to gamma rhythms. (**A-C**) Spike correlates of radiatum slow gamma. (A) An illustrative raw data excerpt from a silicon probe recording shows a radiatum slow gamma event. Individual troughs of slow gamma are demarcated by green triangles. (B) LFP averages triggered by slow gamma troughs (top panel) are accompanied by averages of principal cell (middle panel) and interneuron (bottom panel) activities. These findings are an aggregation of 22 recording days from 10 mice. Within each recording session, the combined activity of either all principal cells or all interneurons were aligned to a single slow gamma trough per theta cycle. Subsequently, an average was computed across all the recording sessions. (C) This panel displays the average gamma phase histogram for both principal cells and interneurons, derived from the same recording days as mentioned in B. This analysis considered only cells that had a significant coupling to the slow gamma (p<0.01), resulting in data from 492 principal cells and 82 interneurons. (**D-F**) Same as in A-C, but for lacunosum-moleculare mid gamma. (E) This panel parallels the analysis in B but pertains to data from recording sessions utilizing a lacunosum-moleculare tetrode. This dataset includes findings from 32 recording days across 13 mice. (F) Same as C but for the mid gamma. From this analysis, 333 principal cells and 70 interneurons were identified with significant coupling to mid gamma oscillations. (**G-I**) Similarly, a sample raw data snippet from silicon probe recordings highlights fast gamma oscillations, with those from the pyramidal layer highlighted in red and from the radiatum in green (G). (H) Same analysis as in B but for pyramidal fast gamma, radiatum fast gamma and ripples, as labeled. For the pyramidal fast gamma and ripples, data from 59 recording days (having both awake and sleep periods) from 20 mice were analyzed. For the radiatum fast gamma data we analyzed data from 17 recording days across 11 mice with a distal radiatum tetrode (Supplementary Figure 5). (I) same analysis as C. From this analysis, 789 principal cells and 220 interneurons were identified with significant coupling to pyramidal fast gamma oscillations, and 112 principal cells and 42 interneurons for radiatum fast gamma.

We subsequently analyzed the modulation of CA1 cell spikes by fast gamma oscillations recorded either from the pyramidal layer or the distal radiatum (**Figure 4G-I**). Using gamma trough-triggered averages for each layer-specific fast gamma, we discerned 2 to 3 cycles of fast gamma modulation in the spiking activity of CA1 principal cells and interneurons (**Figure 4H**; left and middle columns). These findings provide evidence that genuine oscillations at this higher frequency exist in the hippocampus, beyond any potential contamination from spike-leakage artifacts. To draw a comparison, we repeated the same analysis for ripples oscillations, which share a similar frequency range as fast gammas (**Figure 4H**; right column). We found that aligning the spiking activity of both principal cells and interneurons to a single ripple trough per SWR event revealed at least 4 ripple cycles. Regarding the average firing phase, CA1 units consistently preferred the onset of the ascending phase of pyramidal fast gamma troughs, with principal cells notably preceding interneurons (**Figure 4I**; left panel). This temporally structured firing relationship with pyramidal fast gamma closely resembled that observed for ripples (**Figure 4I**; right panel). Conversely, we observed an opposite spike timing relationship in radiatum fast gamma oscillations, with the firing of CA1 interneurons recorded in the pyramidal layer occurring at earlier phase compared to principal cells (**Figure 4I**; mid panel). This pointed to different generating mechanisms for the two CA1 (pyramidal versus radiatum) fast gamma oscillations.

### Profiling deep versus superficial CA1 pyramidal sub-layers

Recent studies have highlighted differences in the firing characteristics of CA1 principal cells within the radial axis of the pyramidal layer (Mizuseki et al., 2011; Navas-Olive et al., 2020; Soltesz and Losonczy, 2018; Valero and de la Prida, 2018). We thus examined whether our feature manifold approach offered the resolution necessary to discern sub-populations of (tetrode-recorded) principal cells within the pyramidal layer. The gradual shift of the sharp-wave waveform recorded along the pyramidal layer radial axis with silicon probes provided the foundation for this estimation. Specifically, the waveform started as a positive deflection near the oriens layer, subtly shifting to a minor negative deflection in the pyramidal layer, before becoming the distinct negative sharp-wave in the radiatum layer (**Figure 5A**). By then projecting each tetrode signal onto the manifold, we obtained a bimodal distribution of estimated depths (**Figure 5B**; left panel). This bimodal organization reflected that even minor adjustments in tetrode positioning around the polarity inversion point of the sharp-wave resulted in pronounced shifts in the manifold feature space, producing a gap in the projection distribution. Consistent with the silicon probe recordings (**Figure 5A**), the tetrodes localized in the pyramidal layer closer to the radiatum layer exhibited a stronger theta-nested slow gamma component than those localized near the oriens layer (**Figure 5B**; right panels). In fact, we found a strong relationship between the amplitude of the sharp-wave and the intensity of theta-nested slow gamma across recording channels (**Supplementary** Figure 6A; Spearman correlation = 0.53, p < 10^-10^), while no such correlation was observed with mid gamma intensity (Spearman correlation = 0.027, p = 0.46).

**Figure 5:**
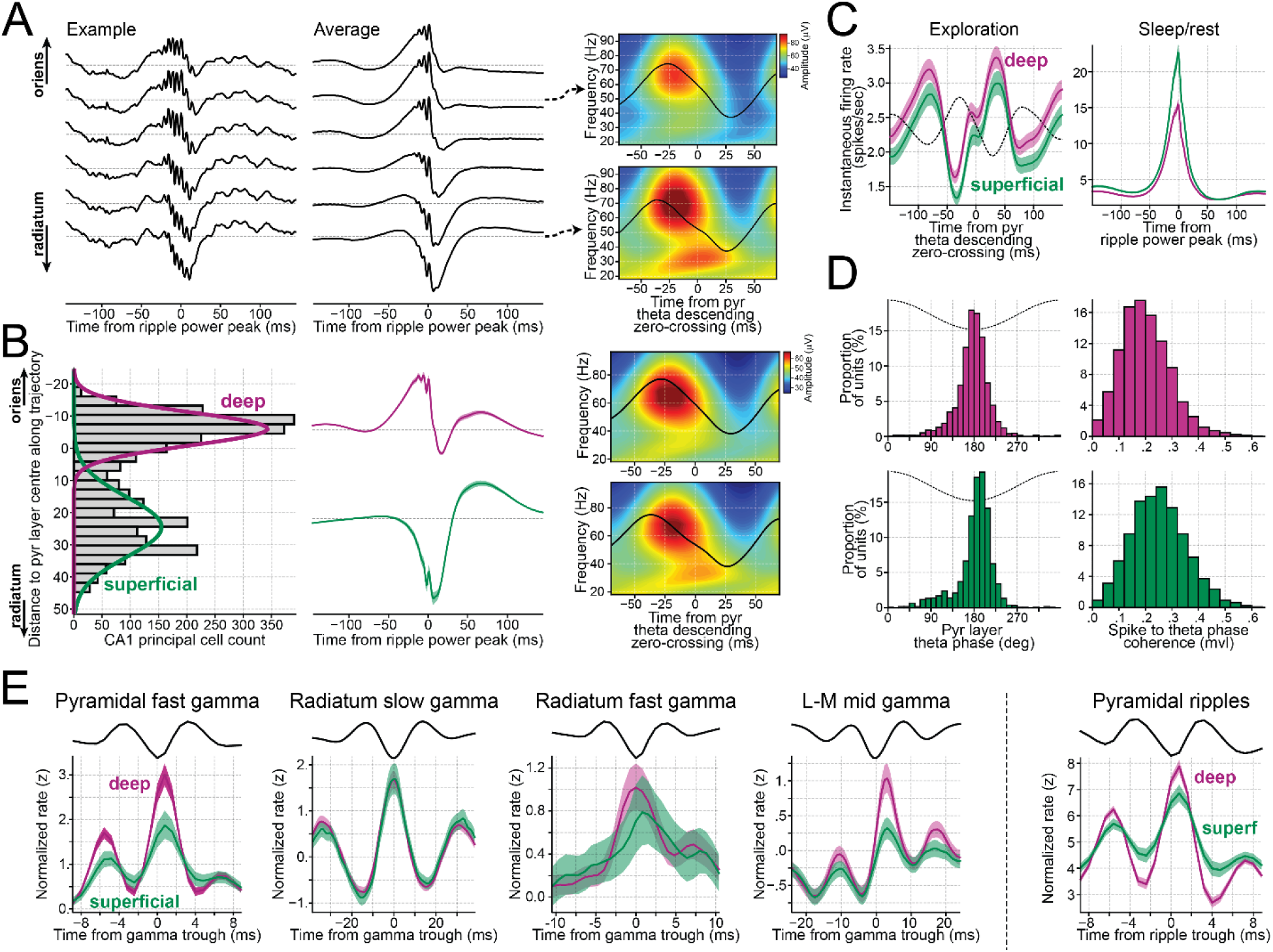
Firing properties of deep and superficial pyramidal sub-layer principal cells. **(A)** Axial profile of SWR waveforms recording from a silicon probe. Left panel shows a single SWR event waveform and mid panel shows the average SWR waveform across all events. The spacing between contacts of this silicon probe is 20μm. The right panels show the amplitude of a range of gamma frequencies (heatmap) and the local LFPs aligned to the pyramidal theta descending zero-crossing for the channels indicated by arrows, calculated in the same manner as in Figure 2. **(B)** Classification of neurons as deep and superficial pyramidal sub-layer principal cells from their projection on the feature manifold. The left panel shows the distribution of the tetrodes of recorded single units projected onto the linearized trajectory introduced in Figure 1E. Overlayed traces denote Gaussian components estimated by a GMM applied to the distribution. Mid panel shows SWR waveforms as recorded from the tetrodes assigned to each Gaussian component. The right panels show the mean theta phase-gamma amplitude profile of the same tetrodes, calculated as in A for the silicon probe channels. **(C)** Mean instantaneous firing rate of deep and superficial pyramidal cells during theta oscillations and SWR. Left panel shows the activity of either deep or superficial pyramidal cells aligned to the pyramidal theta descending zero-crossing, as labelled. The right panel shows a similar analysis but for activity aligned to the power peak of sleep SWRs. (**D**) Theta coupling of deep and superficial pyramidal cells. The left panels show the histogram of the mean firing theta phase for deep and superficial cells. The right panels show the distribution of the coupling level (mean vector length) of these two populations. (**E**) Z-scored instantaneous firing rate of both deep and superficial cells, aligned to the troughs of CA1 gammas and ripples, as indicated. Only one trough is considered per theta cycle. The line atop each panel displays the average LFP recording for the corresponding layer, aligned to the same references. Across all panels, lines represent means, while the shaded areas indicate 99% bootstrap confidence intervals.

Next, we categorized CA1 pyramidal cells into deep and superficial by applying a 2-component Gaussian Mixture Model classifier to their estimated depths (**Figure 5B**). During awake theta oscillations, deep pyramidal sub-layer cells significantly fired at a higher rate (**Figure 5C**; 2.56 spikes/sec, 99% CI: 2.43 – 2.69) compared to superficial sub-layer cells (2.24 spikes/sec, 99% CI: 2.11 – 2.37; p < 10^-5^ for both bootstrap and permutation tests). During SWR events in sleep/rest (analyzed using a 10-ms window centered at the ripple power peak), superficial cells fired significantly more frequently (**Figure 5C**; 19.90 spikes/sec, 99% CI: 18.96 – 20.88) than deep cells (13.62 spikes/sec, 99% CI: 13.13 – 14.14; p < 10^-5^ from both bootstrap and permutation tests). The mean theta firing phases also distinguished the two pyramidal sub-layer cell groups. Specifically, spikes from deep principal cells exhibited an earlier mean theta phase relative to superficial cell spikes (**Figure 5D** and **Table 1**; p < 10^-5^ for both bootstrap and permutation tests). When assessing spike-to-theta phase coherence, superficial cells exhibited a more pronounced coupling than deep cells (**Figure 5D** and **Table 1**; p < 10^-5^ for both bootstrap and permutation tests). This difference in coherence stemmed from deep pyramidal sub-layer cells firing proportionally more spikes around the theta peak compared to superficial cells (**Figure 5C**), nudging their mean phase closer to the peak, and resulting in diminished modulation depth.

**Table 1:**
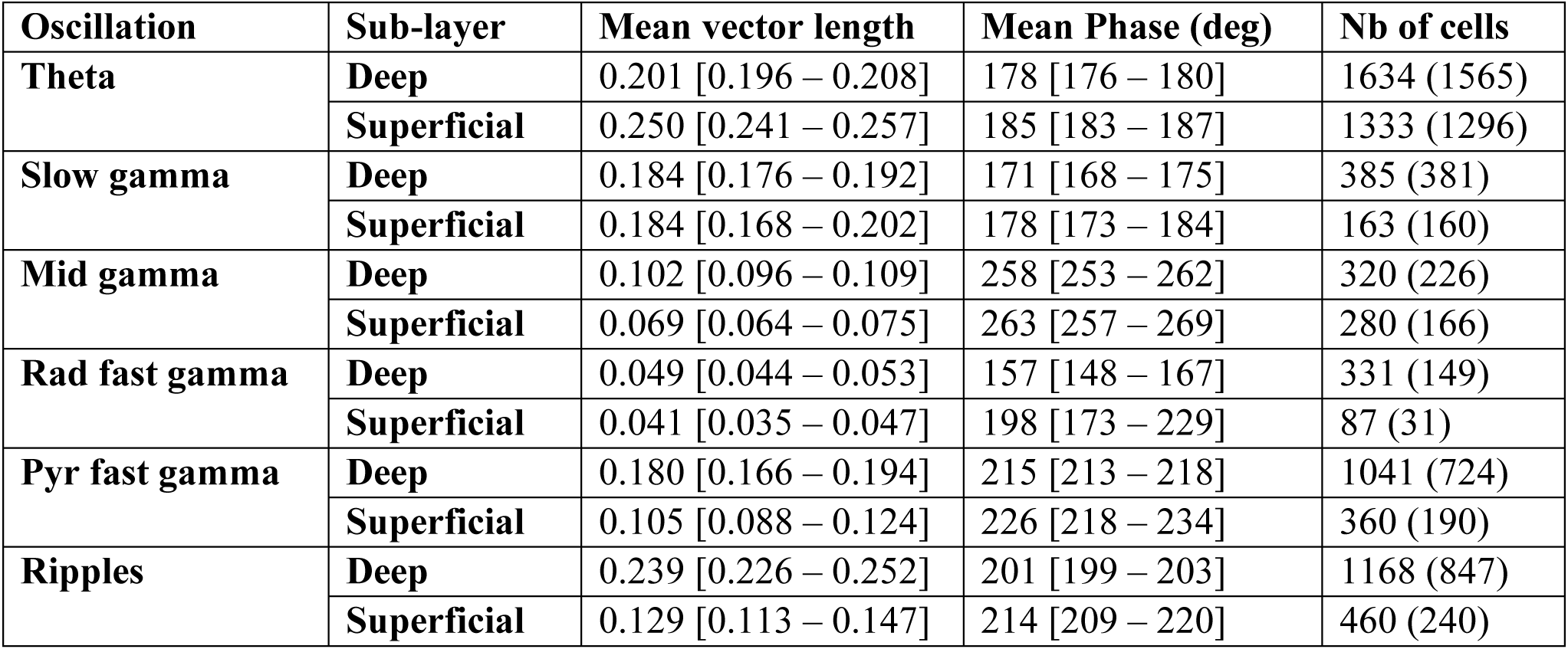
Oscillatory coupling of CA1 deep and superficial pyramidal sub-layer principal cells. The table shows the mean and 99% bootstrap confidence intervals (in square brackets) for mean vector length and mean phase. Analyses include cells with a minimum of 250 spikes for the given oscillation. In addition, only cells with a mean vector length higher than 0.05 and a p value lower than 0.01 were considered for the mean phase analysis. The ’Nb of cells’ column specifies the number of cells included in the mean vector length analysis, with the number for the mean phase analysis indicated in parentheses. Deep cells exhibited significantly stronger coupling to pyramidal fast gamma, mid gamma, radiatum fast gamma, and ripples (p < 10^-5^ for both rate matching and permutation tests in pyramidal fast, mid gamma, and ripples; p = 0.013 and p = 0.0042 for permutation and rate matching tests in radiatum fast gamma, respectively).

| Oscillation | Sub-layer | Mean vector length | Mean Phase (deg) | Nb of cells |
| --- | --- | --- | --- | --- |
| **Theta** | **Deep** | 0.201 [0.196 – 0.208] | 178 [176 – 180] | 1634 (1565) |
|  | **Superficial** | 0.250 [0.241 – 0.257] | 185 [183 – 187] | 1333 (1296) |
| **Slow gamma** | **Deep** | 0.184 [0.176 – 0.192] | 171 [168 – 175] | 385 (381) |
|  | **Superficial** | 0.184 [0.168 – 0.202] | 178 [173 – 184] | 163 (160) |
| **Mid gamma** | **Deep** | 0.102 [0.096 – 0.109] | 258 [253 – 262] | 320 (226) |
|  | **Superficial** | 0.069 [0.064 – 0.075] | 263 [257 – 269] | 280 (166) |
| **Rad fast gamma** | **Deep** | 0.049 [0.044 – 0.053] | 157 [148 – 167] | 331 (149) |
|  | **Superficial** | 0.041 [0.035 – 0.047] | 198 [173 – 229] | 87 (31) |
| **Pyr fast gamma** | **Deep** | 0.180 [0.166 – 0.194] | 215 [213 – 218] | 1041 (724) |
|  | **Superficial** | 0.105 [0.088 – 0.124] | 226 [218 – 234] | 360 (190) |
| **Ripples** | **Deep** | 0.239 [0.226 – 0.252] | 201 [199 – 203] | 1168 (847) |
|  | **Superficial** | 0.129 [0.113 – 0.147] | 214 [209 – 220] | 460 (240) |

Given that low and high firing rate CA1 principal cells are hypothesized to exhibit different properties and functions (Gava et al., 2021; Grosmark and Buzsáki, 2016), it remained plausible that the observed disparities in mean theta phase and coherence in fact arose from firing rate differences rather than anatomical positioning. To clarify this, we performed an additional analysis, subsampling from both deep and superficial cells to match their firing rate distributions (**Supplementary** Figure 6B). This further supported the fact that the variations in mean theta phase and coherence stemmed from differences in CA1 radial location. Specifically, superficial cells had a later mean theta phase than deep cells (mean phase difference = 7.42 degrees; 99% CI: 6.26 – 8.56 degrees, p < 10^-5^). Similarly, superficial cells were more coupled to the theta phase (mean vector length difference = 0.029; 99% CI: 0.026 – 0.033, p < 10^-5^).

We further examined the coupling levels between gamma oscillations and pyramidal sub-layer cell groups (**Figure 5E** and **Table 1**). While slow gamma coupling was equivalent in both groups (**Table 1**), deep cells showed significantly stronger coupling to pyramidal and radiatum fast gammas, as well as to mid gamma and ripple oscillations, compared to superficial cells (**Table 1**). Overall, our results emphasize the distinct firing properties between deep and superficial CA1 pyramidal sub-layers, underscoring the need to factor in the radial organization when analyzing pyramidal cell activity.

## Discussion

### Exploring the hippocampal circuitry via electrophysiological signatures

The hippocampal formation comprises a complex set of interrelated neuronal circuits, each contributing to mnemonic functions. Understanding the cellular operations and substrates of this brain network is essential for understanding how it supports memory formation, storage, and retrieval. By placing electrodes in the different circuits composing this network, we can capture a myriad of electrophysiological patterns, which reflect not only the activation of these individual circuits but also their dynamic interactions. Thus, through the identification and comprehension of these network activity patterns, we gain important insights into the underlying cellular mechanisms of their corresponding circuits and, ultimately, their contribution to memory.

In this context, we provide a detailed characterization of the diverse electrophysiological patterns along the CA1-DG axis. Using the theta phase reversal and the sharp-wave waveform, we crafted a feature manifold that maps the multiple layers of the hippocampal network (**Figure 1**). We observed that these layers present distinct theta-gamma profiles, delineating their unique electrophysiological hallmarks. Importantly, this laminar profiling is consistent across different mice and recording techniques. We propose that such an approach is adaptable to other layered brain regions and different across animal species. For instance, examining the phase reversal of UP and DOWN states and spindles in the neocortex (Girardeau and Lopes-dos-Santos, 2021; Sirota et al., 2003; Sirota and Buzsáki, 2005; Steriade et al., 1993) could reveal nuanced patterns across its layers. Such a comprehensive analysis could enhance our understanding of neocortical cell properties and inherent oscillations. Additionally, examining various neocortical regions, from primary to associative areas, or comparing the same network across species might provide further insights.

### Network layering and cellular basis of theta-nested hippocampal gamma oscillations

CA1 slow gamma is thought to originate in the radiatum (Belluscio et al., 2012; Lasztóczi and Klausberger, 2016; Lopes-dos-Santos et al., 2018; Schomburg et al., 2014), a layer where CA3 projects to CA1 via Schaffer collaterals (Ishizuka et al., 1990; Witter et al., 2000). This is consistent with observations that CA3 LFPs prominently expresses slow gamma (Schomburg et al., 2014; Tort et al., 2009), and that there is a notable coherence between CA1 and CA3 LFPs within this frequency band (Colgin et al., 2009; Kemere et al., 2013; Schomburg et al., 2014). When CA3 terminals in CA1 are optogenetically suppressed, slow gamma power is reduced while mid gamma oscillations remain unaffected (El-Gaby et al., 2021). Moreover, activating dorsal CA3 parvalbumin-expressing interneurons, which curtails CA3 principal cell spiking output, also leads to a significant decrease in slow gamma oscillations (Aery Jones et al., 2021; López-Madrona et al., 2020). In our study, we highlight the pronounced presence of slow gamma oscillations in the radiatum layer of CA1. While these oscillations can be detected in the pyramidal layer, they are mainly evident in the pyramidal sub-layer proximal to the radiatum, corresponding to channels that display a negative sharp-wave. The slow gamma observed in the pyramidal layer is presumably propagated through volume conduction from the radiatum layer. The dense cellular structure of the pyramidal layer likely acts as a physical barrier, constraining the spread of this oscillation towards the oriens layer. In the lacunosum-moleculare layer, though slow gamma can be observed, it is often overshadowed within its average theta cycle by the more dominant mid gamma oscillations in that layer.

Mid gamma oscillations are proposed to originate from the stratum lacunosum-moleculare (Lasztóczi and Klausberger, 2016; Lopes-dos-Santos et al., 2018; Schomburg et al., 2014), which is the main CA1 target of entorhinal cortex layer 3 inputs (Desmond et al., 1994; Van Groen et al., 2003). Intriguingly, we detected mid gamma activity in more remote layers like the oriens. While volume conduction might account for the detection of oscillations far from their source, slow gamma oscillations emanating from the radiatum layer are not clearly seen in the other side of the pyramidal layer. In addition, despite the potential low pass filtering of currents from the apical dendrites of pyramidal cells as they travel to the soma, the spiking of these cells was still clearly modulated by this rhythm (**Figure 4E**). This probably arises from mid gamma-paced inputs from the entorhinal cortex to CA1 activating interneurons in the radiatum and lacunosum-moleculare layers (Lasztóczi and Klausberger, 2014). By resonating with the mid gamma rhythm, a particular set of specialized interneurons could then modulate the firing patterns of pyramidal cells, fostering synchronicity between their activity and mid gamma-associated inputs. While these interneurons play a pivotal role, it is also worth noting that other cell types may have a different impact. For instance, a recent study suggested that neurogliaform cells in lacunosum-moleculare can decouple cortical gamma inputs and CA1 pyramidal cell firing (Sakalar et al., 2022).

CA1 fast gamma oscillations are primarily contained within the CA1 pyramidal layer, likely reflecting interactions between pyramidal cells and local interneurons (Lasztóczi and Klausberger, 2016, 2014). Although concerns regarding contamination from spike waveform leakage have been substantiated (Belluscio et al., 2012; Scheffer-Teixeira et al., 2013), our prior (Lopes-dos-Santos et al., 2018) and current observations (**Figure 4H**) demonstrate genuine rhythmicity, as observed when aligning spike trains of CA1 neurons to a single fast gamma trough per theta cycle. This underscores that despite potential spike-leakage contamination (Lopes-dos-Santos et al., 2018), pyramidal fast gamma oscillations remain authentic neural rhythms. We further provide findings suggesting that CA1 pyramidal fast gamma oscillations could arise from a pyramidal-interneuron network gamma (PING) circuit (Wang, 2010; Whittington et al., 2000). Within this framework, pyramidal cells activate local interneurons, which then inhibit the activity of the pyramidal cell population through lateral inhibition. Once the inhibition wears off, and if the excitatory drive into the pyramidal cells persists, these cells re-initiate firing, thus beginning a new cycle. We note that this model could similarly account for the ripple oscillations within the same neuronal population (**Figure 4H**). We therefore propose that the most parsimonious explanation is that CA1 pyramidal fast gamma and ripples arise from the same cellular foundation. The distinguishing factor would be the initiating drive: pyramidal fast gamma would be triggered by a theta-associated drive, while ripples are spurred by sharp-wave related inputs. Additionally, optogenetic stimulation of CA1 pyramidal cells can activate this oscillator (Stark et al., 2014). Interestingly, the timing relationship between pyramidal cells and interneurons associated to the radiatum fast gamma (**Figure 4H**) cannot be explained by the same mechanism. Instead, this rhythm likely stems from feedforward inputs directed to the radiatum and/or lacunosum-moleculare layers.

Fast gamma oscillations in the DG have been documented in previous studies (Fernández-Ruiz et al., 2021; Lasztóczi and Klausberger, 2017; Scheffer-Teixeira et al., 2012) and have been identified as distinct from CA1 fast gamma (Lasztóczi and Klausberger, 2017). In our current work, we build from these findings by distinguishing two fast gamma oscillations within the DG. One is more pronounced in the molecular layer, while the other is dominant in the granular layer (**Figure 2**). Earlier studies also identified slow gamma oscillations in the DG (Fernández-Ruiz et al., 2021; Lasztóczi and Klausberger, 2017). Given their theta phase relationship (mirroring CA1 radiatum slow gamma), and the pronounced entrainment of DG cells by CA1 slow gamma oscillations (even more so than CA1 cells), it is postulated that both might share a common oscillatory source (Lasztóczi and Klausberger, 2017). One possibility is that these oscillations initiate in the DG, are transmitted to CA3, and subsequently manifest in CA1 radiatum (Bragin et al., 1995a; Hsiao et al., 2016; Lasztóczi and Klausberger, 2017). Alternatively, a potential source of slow gamma oscillations might be outside the hippocampus, possibly within the medial septum (Király et al., 2023). Moreover, Fernandez-Ruiz et al (2021) recently suggested that DG slow gamma might originate from inputs of the lateral entorhinal cortex layer 2. As we continue our investigations into these oscillations, the importance of discerning their origins and interactions becomes apparent.

### Radial organization of principal cells within the CA1 pyramidal layer

By leveraging our feature manifold, we identified sub-layers within the CA1 *stratum pyramidale* and examined the firing properties of pyramidal cells in freely moving mice using independently moveable tetrodes. We replicated findings from prior studies using silicon probe recordings, thereby possessing a ground truth for the relative positioning of recorded cells. Specifically, we observed that during awake periods of exploratory behavior, CA1 principal cells in the deep pyramidal sub-layer are more active in theta oscillations (Mizuseki et al., 2011), whereas those in the superficial sub-layer are more active in SWRs (Stark et al., 2014). The rate difference in SWRs aligns with *in vivo* patch-clamp studies which indicate that superficial cells experience a heightened excitatory drive during SWR events (Valero et al., 2017, 2015). These consistent observations further validate our LFP-based depth estimation approach. Moreover, we observed that superficial pyramidal sub-layer cells have a more pronounced coupling to the theta phase than their deep counterparts. Conversely, deep cells demonstrated an average heightened synchrony with both ripples and gamma oscillations. Notably, the sole exception was slow gamma, where both principal cell subpopulations exhibited comparable synchrony. These firing properties likely arise from the fact that local fast spiking interneurons exert stronger control in deep principal cells than in superficial ones (Lee et al., 2014). The heightened response of superficial cells to CA3 inputs (Valero et al., 2015) might account for their equivalent modulation with deep cells for slow gamma. It is also possible that deep cells are more responsive to gamma-associated currents. This pronounced response to gamma currents may divert deep cells from the primary sway of theta rhythms, leading them to discharge proportionally more spikes outside their preferred theta phase. Crucially, these findings are not attributable to rate differences between the two principal cell subpopulations. Thus, they are more likely tied to cell radial location within distinct sub-layers, rather than a simple disparity between highly active versus less active cells. The distinction between deep and superficial cells within the CA1 pyramidal layer is now emerging as a critical factor in hippocampal processing (Soltesz and Losonczy, 2018). Our findings underscore that the radial organization of the CA1 pyramidal layer is pivotal for a comprehensive and nuanced appreciation of hippocampal function.

### On the significance of neural oscillations

LFP signals predominantly reflect synaptic activity (Buzsáki et al., 2012). For example, in the SWR complex, the sharp-wave in the radiatum is believed to reflect synchronized excitatory post-synaptic potentials from CA3 ensembles arriving via Schaffer collaterals (Buzsáki, 2015, 1986; Buzsáki et al., 1992). When excitatory or inhibitory post-synaptic potentials occur at a regular pace, they manifest in LFP signals as rhythmic waves, as is the case for gamma oscillations (Buzsáki and Wang, 2012). Different gamma oscillations can be distinguished by the circuit mechanisms driving their associated rhythmic post-synaptic potentials.

Here, we identified four fast gamma oscillations with overlapping frequency bands. These rhythms manifest in distinct layers of the CA1-DG axis and relate differently to the theta cycle. Particularly, the two fast gamma oscillations in CA1 — one in the pyramidal layer and the other deep in radiatum — reveal different timing relationships between pyramidal cells and interneurons. We also observed a beta component, which sits within the slower range of the broad gamma band. Past studies have defined slow (or low) gamma as the oscillation associated with CA3-to-CA1 inputs, most prominent during the descending phase of theta and linked to the CA1 radiatum layer (Belluscio et al., 2012; Bragin et al., 1995; Colgin et al., 2009; Fernández-Ruiz et al., 2017; Lasztóczi & Klausberger, 2016; Lopes-dos-Santos et al., 2018; Schomburg et al., 2014). In contrast, we observed that the beta component arises near the theta peak, being clearer in the DG mid molecular layer (**Supplementary** Figures 3,4). The differences between these patterns (beta versus slow gamma) support the need to identify network oscillations based on their functional and anatomical foundations rather than solely their frequency bands.

In line with this rationale, the present study supports the recent call for a revision in the nomenclature of gamma oscillations to benefit the field of brain network physiology (Fernandez-Ruiz et al., 2023). For example, hippocampal CA1 “slow gamma” might be more aptly termed “CA3-to-CA1 gamma”, thereby emphasizing its anatomical and functional basis over its frequency band. Similarly, CA1 pyramidal layer “fast gamma” would be more appropriately named CA1 “perisomatic gamma” (Lasztóczi and Klausberger, 2014). Such a framework implies that if a hippocampal gamma oscillation in another species has the same cellular foundation, it would then be recognized as the same oscillation even if it has a different frequency band. Conversely, a signal with a similar frequency band but recorded in another brain region, such as the visual cortex, would not be regarded as equivalent. Adopting this circuit-based approach would minimize potential confusions while emphasizing the physiological significance of brain rhythms.

## Acknowledgments

We would like to thank S. McHugh and R.M.M. Santiago for commenting on a previous version of the manuscript; B. Micklem for technical assistance; all members of the Dupret lab for feedback during the project. This work was supported by the Biotechnology and Biological Sciences Research Council UK (awards BB/S007741/1 and BB/N002547/1) and the Medical Research Council UK (programme MC_UU_00003/4 and award MR/W004860/1). D.B. is supported by a Biotechnology and Biological Sciences Research Council (BBSRC) UK studentship (BB/T008784/1) and a Scatcherd European Scholarship.

## Methods

### Animals

These experiments used 32 adult mice (4–6 months old; see Supplementary Table S1). Animals were housed with their littermates up until the start of the experiment. All mice held in IVCs, with wooden chew stick, nestlets and free access to water and food *ad libitum* in a dedicated housing facility with a 12/12 h light/dark cycle (lights on at 07:00), 19–23°C ambient temperature and 40–70% humidity. Experimental procedures performed on mice in accordance with the Animals (Scientific Procedures) Act, 1986 (United Kingdom), with final ethical review by the Animals in Science Regulation Unit of the UK Home Office.

### Surgical procedure

All surgical procedures were performed under deep anesthesia using isoflurane (0.5–2%) and oxygen (2 l/min), with analgesia provided before (0.1 mg/kg vetergesic) and after (5 mg/kg metacam) surgery.

For silicon probe recordings, mice were implanted with a single-shank silicon probe (Supplementary Table 1) under stereotaxic control in reference to bregma, using central coordinates -2.0 mm anteroposterior from bregma, +1.7 mm lateral from bregma, and an initial depth of 1.5 mm ventral from the brain surface to span the somato-dendritic axis of CA1 principal cells and reach the DG. Following the implantation, the exposed parts of the silicon probe were covered with Vaseline® Healing Jelly, after which its plastic drive was secured to the skull using dental cement and stainless-steel anchor screws inserted into the skull. Two of the anchor screws, both above the cerebellum, were attached to a 50 µm tungsten wire (California Fine Wire) and served as ground. For the recordings, the silicon probe was positioned along the CA1-to-DG axis, using the rotations applied to its holding screw.

For tetrode recordings, mice were similarly implanted with a single microdrive containing 14 independently movable tetrodes, targeting the *stratum pyramidale* of the dorsal CA1 hippocampus. Tetrodes were constructed by twisting together four insulated tungsten wires (12 μm diameter, California Fine Wire) which were briefly heated to bind them together into a single bundle. Each tetrode was loaded in one cannula attached to a 6 mm long M1.0 screw to enable its independent manipulation of depth. The drive was implanted under stereotaxic control in reference to bregma using the following coordinates. For pyramidal layer channels, the span was between AP -1.4 to –2.7 mm and ML 0.9 to 2.4 mm. Tetrodes that delved deeper than the pyramidal layer were positioned within the range of AP –1.9 to –2.5 mm and ML 0.9 to 1.7 mm. The initial depth of the tetrodes during the implantation surgery was 1.0 mm ventral from the brain surface. The distance between neighboring tetrodes was 350 μm. Following the implantation, the exposed parts of the tetrodes were covered with paraffin wax, after which the drive was secured to the skull using dental cement and stainless-steel anchor screws inserted into the skull. Two of the anchor screws, both above the cerebellum, were attached to a 50 µm tungsten wire (California Fine Wire) and served as ground. For the recordings, each tetrode was lowered along the vertical axis to reach the CA1 pyramidale layer, using the rotations applied to its tetrode cannula-holding screw and the electrophysiological profile of the local field potentials in the hippocampal ripple frequency band, with final depth position subsequently confirmed by histology of anatomical tracks.

### Recording procedures

Following full recovery from the surgery, each mouse was first handled in a dedicated handling cloth, connected to the recording system, and exposed to an open-field enclosure to be familiarized with the recording procedure over a period of one week prior to the start of the experiment itself. During this period, animals were habituated to a sleep box (outer dimension: 12 cm width; 16 cm height) containing bedding from their home cage. This sleep box served for all sleep recordings. When conducting awake sessions, the mice were set in open-field enclosures that varied in shape and had maximum side dimensions of 46 cm. For recordings using silicon probes, the position of the silicon probes was gradually adjusted along the dorsal-ventral axis until the pyramidal cell layer SWR events were recorded by uppermost region of the silicon probe. Once positioned, the probes remained stable across various recording sessions. For tetrode recordings, the tetrodes were individually moved from their original post-surgery location to the CA1 pyramidal layer, which could be distinctly identified by the pronounced presence of SWR events. Before beginning the recordings each day, the position of the tetrodes was fine-tuned to optimize both the clarity and the number of spike waveforms, based on visual assessment. Moreover, tetrodes targeting layers ventral from the CA1 pyramidal layer, or the dentate gyrus were moved downwards until they reached their intended target (see *Tetrode feature-based placement* section for details). Once the tetrodes were positioned for that recording day, we allowed for a 90-minute break before commencing the first recording session, ensuring sufficient time for the tissue to adjust. As a day of recording concluded, the tetrodes located in the pyramidal layer were cautiously retracted by about 125μm in the direction of the stratum oriens. This preventive step ensured that the pyramidal layer remained unharmed overnight.

### Multichannel data acquisition and position tracking

The extracellular signals from each recording channel were amplified, multiplexed, and digitized using a single integrated circuit located on the head of the animal (RHD2164, Intan Technologies; http://intantech.com/products_RHD2000.html; pass band 0.09 Hz to 7.60 kHz). The amplified and filtered electrophysiological signals were digitized at 20 kHz and saved to disk along with the synchronization signals (transistor-transistor logic digital pulses) reporting the animal’s position tracking. The location of the animal was tracked using three differently colored LED clusters attached to the electrode casing and captured at 39 frames per second by an overhead color camera (https://github.com/kevin-allen/positrack/wiki).

### Local field potential signals

LFP signals were processed by first applying an anti-aliasing filter (8^th^-order Chebyshev type I filter) to the wide band signals sampled at 20kHz. These signals were then downsampled to 1,250Hz using the decimate function from the signal submodule of Scipy (version 1.11.2).

### Detection of SWR events

For SWR event detection, the LFPs were initially referenced against a channel where CA1 ripples were not observed. The resulting differential signal was filtered through a ripple band filter (80-250Hz, 4^th^-order Butterworth filter) and through a control high-frequency band filter (200-500Hz, 4^th^-order Butterworth filter). For all ripple analyses, we used the Hilbert transform to compute instantaneous envelopes and phases.

Candidate SWR events were demarcated when peaks of the ripple band envelope surpassed a threshold set at fivefold its overall median value. When multiple peaks arised within a 20-ms timeframe, only the highest peak was kept. The boundaries of each candidate event, onset and offset, were pinpointed where the envelope dipped below half the detection threshold.

We further calculated the number of ripple cycles within each event. This was achieved by determining the phase change across the event, which is the difference between the unwrapped phase at the offset and the onset phase (for example, 1800 unwrapped phase difference from onset to offset equals to five cycles as 1800/360=5). The event’s mean frequency was then derived by dividing the number of its cycles by its duration in seconds (for example, 6.75 cycles in 50ms would yield a 135Hz mean frequency event). Finally, each candidate event underwent a series of checks: (1) The ripple band power (derived from squaring the mean ripple amplitude) in the detection channel should exceed twice the magnitude obtained for the reference channel. This confirms that the detected events in the differential signal had a stronger presence in the detection channel; (2) The mean frequency of the event should surpass 100 Hz; (3) The event must comprise a minimum of 4 complete ripple cycles; (4) The power in the ripple band should be at least double compared to the control high frequency band.

### Determination of the reference CA1 pyramidal layer channel

For recording, we used a single reference channel for SWR events and theta oscillations, which was defined as the channel having the highest ripple band score. This was defined as the power in the ripple band (100-250Hz) divided by the power within a surrounding frequency band (70-300Hz). The power in each band was calculated by means of a Welch spectrum with 4-second Hann windows with 50% overlap.

### Extraction of theta oscillations from LFPs

To isolate theta oscillations from the LFP data, we employed the masked Empirical Mode Decomposition method (Deering and Kaiser, 2005; Huang et al., 1998) as implemented by Quinn et al (2021a). For optimal results, particularly with CA1 recordings, we adopted the mask sift procedure with specific mask frequencies set at 350, 200, 70, 40, 30, and 7 Hz, following the parameters optimized in (Quinn et al., 2021b) grounded in (Fosso and Molinas, 2018). For each mask, the amplitude was set to three times the standard deviation of the input signal. This procedure decomposes each LFP signal into oscillatory components termed intrinsic mode functions (IMFs) from faster to lower frequency components. Upon completion of this procedure with the beforementioned parameters, six IMFs and a residue were computed, with IMF-6 effectively isolating theta oscillations.

To delineate individual theta cycles, we began by pinpointing peaks and troughs—i.e., the local maxima and minima, respectively—of the theta IMF derived as previously outlined. The residue of the LFP not captured by the first six IMFs were defined as the lower frequency component of the signal and its envelope was used as amplitude threshold for retaining peaks and troughs for the next step. We next defined each peak-trough-peak sequence as a candidate theta cycle. We took as a valid cycle sequences having their peak-trough and trough-peak intervals falling within the 31 to 100 ms range (corresponding to the half period of cycles with frequencies ranging from ∼16 to 4 Hz); and peak-to-peak distance was between 71 ms (equivalent to ∼14 Hz) and 200 ms (equivalent to 5 Hz).

For each validated cycle we found six control points: the zero-crossing prior to the first peak, the peak itself, the subsequent zero-crossing post the first peak, the trough, and the zero-crossing following the trough. Then, we computed the instantaneous theta phase for each timestamp through a linear interpolation of the control points (Belluscio et al. 2012; Lopes-dos-Santos et al. 2018).

### Current source density analysis

In this study, we applied CSD analysis (Brankack et al., 1993; Mitzdorf, 1985) to the event-triggered averages from LFP recordings captured via linear silicon probes. We calculated these averages by centering intervals of the LFP signals around event timestamps. For instance, when focusing on theta, we centered our averages around the descending zero-crossings of detected theta cycles in a recording. After establishing these averages for specific events, the current source density signal at channel *n* and a given time point was computed as:

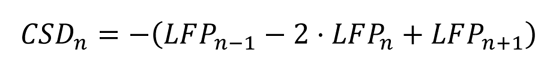

where, *n−1* and *n+1* refer to the channels immediately above and below *n*.

To standardize the spatial resolution of CSDs across silicon probes with varying channel spacings, we applied a Gaussian kernel smoothing with a standard deviation of 50 μm.

### Dentate spikes detection and classification

Dentate spikes in silicon probe recordings were predominantly identified during sleep/rest periods, given their frequent occurrence in these brain states (Bragin et al., 1995b; Lensu et al., 2019). Initially, LFPs from the dentate gyrus region were subtracted using a reference channel located at the top of the probe, typically positioned in the cortex or high in the oriens/alveus region. After this subtraction, the differential signal underwent filtering within a frequency range of 1-200 Hz using a 4^th^-order Butterworth filter. Peaks in this filtered differential signal that surpassed a threshold, set at seven times the median absolute value, were identified as candidate dentate spike events. To ensure the removal of high-amplitude artifacts, any candidates with peaks exceeding a threshold—defined as the 75^th^ percentile of the peak distribution added to 20 times its interquartile range—were excluded. This detection procedure was applied to channels within the dentate gyrus region, determined through visual inspection, and events from these channels were then combined. If multiple events were detected within a 50ms window, only the event with the highest peak was retained. To categorize dentate spikes into DS1 and DS2 types, we analyzed the CSDs of each event, computing the CSD at its peak from the pyramidal layer channel to the final dentate gyrus channel. We then employed Principal Component Analysis on these CSD vectors, projecting each individual CSD onto the first principal component. Classification was then achieved using a 2-component Gaussian Mixture Model (GMM). In recordings that covered the molecular and granular layers of the dentate gyrus, this method consistently identified two distinct CSD patterns, each with primary sinks located at different points within the dentate gyrus channels. The DS1 group was characterized by its primary sink situated closer to CA1, while the DS2 group had its main sink nearer to the hilus. If the GMM-defined classes showcased identical primary sinks, we labeled all dentate spike events as DS1. This designation indicated that the recordings lacked a granular layer channel.

### LFP feature manifold

We used the silicon probe recording dataset to construct the LFP feature manifold. For each mouse, we computed the sharp-wave and theta waveforms from each recording channel, thus extending from the CA1 pyramidal layer to the DG granular layer (**Figure S2A**). These waveforms represent the average raw LFP signals for each of these individual channels, centered around either the ripple power peaks or the theta descending zero-crossings. The length of these triggered averages was 500 ms for sharp-waves and 150 ms for theta. For uniformity across mice, we sampled channels at 50 μm steps (40 μm for mice with 20 μm-spaced channels; see **Table S1**). Each waveform was then z-scored individually to focus this analysis on the waveform shapes and not the relative amplitudes across channels. We next extracted the first four principal components from these waveforms (**Figure S2B**). Using these principal component projections, the manifold was then computed employing the Isomap method from the manifold submodule of sklearn (version 1.3; with parameters n_neighbors = 15 and n_components = 2). We next obtained a trajectory that connects layer centers as follows. To enhance resolution between distantly situated clusters in the feature space (e.g., between radiatum and lacunosum-moleculare), we incorporated intermediate points prior to interpolation. The number of intermediate points between successive layer pairs was determined by dividing the distance between points in the 2-D feature space by 20. With this derived count, we selected an equivalent number of equidistant channels in the real anatomical space for each mouse. Subsequently, we calculated the average for each control point (layer coordinates and intermediate points) across all the mice. The final trajectory (Figure 1E, black trace; and **Figure S2C**, right panel) was determined by quadratic interpolating these average control points.

### Classification of layers from manifold projections

We evaluated the separability of the layers within the feature manifold using cross-validated classification. Leveraging a leave-one-out methodology, each classifier was trained on all available data points except one, which was subsequently used for testing. It is worth noting that each mouse contributed a single layer channel; hence, the classifiers primarily decoded the layer data based on datasets from other mice. We utilized both the k-neighbors classifier (parameters: weights=‘uniform’, algorithm=‘auto’, and k=4) and the nearest centroid classifier (using metric=‘euclidean’ and shrink_threshold=None). Both classifiers were obtained from the ‘neighbors’ submodule of sklearn (version 1.3.0). The choice of k=4 for the k-neighbors classifier corresponds to the smallest sample size across layers.

### Supra-theta wavelet spectrograms

Spectrograms were generated using the complex Morlet Wavelet Transform technique. For this analysis, we selected a range of 24 log-spaced frequencies, spanning from 18 Hz to 310 Hz unless stated otherwise. We applied L1 normalization to each wavelet kernel. In L1 normalization, the sum of the absolute values of the elements in a vector is ensured to equal 1. This ensures that the wavelet does not amplify or attenuate the amplitude of individual frequency components.

### Gamma filtering and instantaneous phase and amplitude

For amplitude or phase analyses of specific gamma oscillations in this study, we filtered the LFP signals within the relevant bands using a 4^th^ order Butterworth filter. The bands were defined by the following cutoff frequencies: 20-45Hz for slow gamma, 50-100Hz for mid gamma, and 100-250Hz for fast gammas. We then determined the instantaneous amplitude and phase using the Hilbert transform.

### ICA-based extraction of fast gamma oscillations

Isolating fast gamma oscillations poses a challenge due to their relatively low amplitude, especially in tetrodes that exhibit higher amplitude mid gamma oscillations. Any overlap with mid gamma oscillations can contaminate the signal filtered for the fast gamma range, given the comparable magnitude of the high-frequency tail of the mid gamma spectrum and the fast gamma oscillation itself. To address this, we adopted an ICA-based method inspired by Fernández-Ruiz & Herreras (2013). This approach leverages spatial data from silicon probe recordings to differentiate fast gamma from mid gamma oscillations. Specifically, ICA was applied to LFP traces extending 200 μm around a targeted channel with fast gamma interest (e.g., pyramidal, distal radiatum, mid moleculare, or granular layer), and we extracted independent components with peak frequencies above 100Hz. We used the amplitude of such signals for the fast gammas in Figure 2B.

### Determination of gamma main frequencies

To discern the primary frequency components of each gamma oscillation, we first pinpointed the strongest bursts within the theta cycles by computing the envelope peaks for the appropriate gamma band during individual cycles, retaining only the top quartile of these peaks. Using these peaks, we generated the mean spectra by averaging wavelet spectrograms within a 40-ms window centered on the burst peaks. Before averaging across mice (Figure 2C), we normalized the triggered average spectrogram of each individual mice for its overall standard deviation.

### Tetrode feature-based placement

To target layers below pyramidal CA1 with tetrodes, we implemented a stepwise, gradual lowering approach. We recorded short sessions during both sleep and wakefulness (usually ranging from 5 to 10 minutes) to project LFP features onto the feature manifold (as depicted in Figure 1D**,E**, and **Supplementary** Figure 2). We ensured that tetrodes were adjusted by no more than ∼60μm in a 20-minute interval. Once a tetrode reached its target layer, we allowed a 90-min break before starting the recordings.

Regarding the tetrode data analysis on spike correlations with specific gamma oscillations, we performed the analysis within the appropriate layer for the given rhythm (Figure 4 and **5**). Slow gamma was consistently assessed from radiatum channels, while mid gamma was derived from lacunosum-moleculare channels. In contrast, fast gammas were measured from their respective local layers.

### Spike detection and unit isolation

Spike sorting and unit isolation were performed with an automated clustering pipeline using Kilosort via the SpikeForest framework (Magland et al., 2020; Pachitariu et al., 2016). To KiloSort data acquired using tetrodes, the algorithm restricted templates to channels within a given tetrode bundle, while masking all other recording channels. The resulting clusters were verified by the operator using cross-channel spike waveforms, auto-correlation histograms, and cross-correlation histograms. Each unit used for analyses showed throughout the entire recording day stable spike waveforms.

### Principal cell versus interneuron classification

To evaluate the waveform consistency for each unit, we focused on the waveform with the maximum amplitude across the tetrode channels for each cluster. Our primary aim was to compare the prominence of a unit mean waveform amplitude to the standard deviation stemming from all its spikes. This waveform score was defined as:

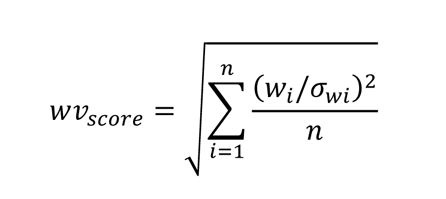

where *w_i_* is the value of the mean waveform at sample *I*, *σ_i_* is the standard deviation at sample *i* across all spikes, and n is the number of waveform samples (in this context, n=32). This metric essentially quantifies the relative magnitude of the mean waveform amplitude against the spike-to-spike variability. Clusters presenting a waveform score below 0.75 or spike trains with a refractory period violation exceeding 2% (quantified as the proportion of intervals shorter than 2 ms in the ISI distribution) were categorized as multi-units and excluded from subsequent analyses. Additionally, clusters displaying positive spikes were also disregarded, as these could potentially originate from non-somatic spikes (Someck et al., 2023). Only the units that passed these criteria were considered well isolated and included in further analyses.

Finally, we categorized units as either putative interneurons or pyramidal cells, relying on the width of their waveform as indicated by the trough-to-peak latency. To achieve a finer resolution, we increased the 32 waveform points sampled at 20kHz by a factor of 100, utilizing quadratic interpolation (employing the interpolate function from scipy). In a prior dataset of ∼4,000 well-isolated neurons, we noted a bimodal distribution in trough-to-peak latency. Fitting this with a 1-dimensional, 2-component Gaussian Mixture Model (GMM) from sklearn, we set the classification threshold where the two Gaussian components intersect. Using this threshold on our current dataset, units with latencies above were labeled as putative pyramidal cells, and those below as putative interneurons. In total this study included 3,012 principal cells and 546 interneurons.

### Gamma and ripple trough triggered averages

In Figures 4B, E, H, and 5E, we computed the mean spiking activity or LFP signals centered around the most prominent troughs of the targeted gamma rhythm or ripples. We began by filtering the LFP signals from the specific layer within the relevant frequency range as mentioned above. For gamma oscillations, the most profound trough within each theta cycle was identified and we used only the top quartile in terms of amplitude for the triggered-average analyses. By aligning data (spike trains or LFP signals) to one distinct gamma trough within each theta cycle, we can accurately depict the gamma rhythmicity. This method gives clarity on the existence and sequence of following gamma cycles, all referenced from that single trough. Similarly, for ripple triggered averages, we filtered pyramidal LFPs within 100-250Hz and used the deepest trough of ripples for each SWR event.

### Spike to phase coherence analysis

Theta oscillations are asymmetrical (non-linear), resulting in an uneven distribution of phases. In recordings from the pyramidal layer, the rising phase is shorter than the falling phase, leading to a predominance of the latter in the recordings. This disparity can skew coherence analyses, showing an artificial increase in spikes associated with the longer, falling phase of the theta oscillations. To correct this, we normalized the spike-phase coherence against the likelihood of a spike occurring in each theta phase bin. This was done by dividing the distribution of theta phases associated with neuron spikes by the overarching theta phase distribution. These distributions were represented as histograms with 64 equally spaced phase bins. The spike-to-phase coherence was quantified using the mean vector length, with each phase bin center weighted by its assigned probability. This approach was also applied to ripples. For gamma phase coherence, the methodology mirrored the theta analysis, but with a modification due to the transient nature of gamma oscillations. We introduced a threshold based on the gamma envelope, considering only the spikes and gamma phases where the gamma envelope surpassed its 75^th^ percentile. This precaution ensured our data was sourced from periods with a genuine gamma presence in the signal.

For a given oscillation, we restricted phase coherence analysis to units that recorded at least 250 spikes coincident with that oscillation, following the criteria outlined above. To assess its significance, we employed a spike shift control. For each neuron, we shifted spikes to random time points, ensuring the original theta phase and gamma amplitude were preserved (Lopes-dos-Santos et al., 2018). We repeated this surrogate process to obtain a null hypothesis distribution with 10,000 surrogate values. The p-value for each neuron was then calculated based on the proportion of surrogate values that matched or exceeded the actual coherence value, indicating the likelihood of such an observation occurring by chance. Only units demonstrating significant coupling (p<0.01) had their mean phases estimated.

### Classification of deep and superficial principal cells

In the analyses depicted in Figure 5, we distinguished principal cells as either deep or superficial based on the LFP features of the tetrodes from which neuronal activity was recorded. The categorization relied on the linearized projection of these features to the feature manifold (refer to Figure 1E, right). Notably, the distribution of these projections across our dataset was distinctly bimodal. To model this, we applied a 1-dimensional, 2-component Gaussian Mixture Model (GMM) from sklearn. The resulting Gaussians are illustrated in Figure 5B superimposed on the projection distribution. We set the classification threshold for deep versus superficial at the intersection of these Gaussian distributions, which approximately equated to 5.5.

### Bootstraps and permutation tests

For analyses in Figures 4 and 5, we used permutation tests to determine the significance of differences between cell groups. The core of this method involves comparing the actual mean difference between two groups to a distribution of mean differences generated from random group assignments. As an illustration, when assessing the mean rate difference between deep and superficial cells, we randomly shuffled cell labels and stored the mean difference of these shuffled groupings. This shuffling was done 100,000 times to produce a null hypothesis distribution, representing the likelihood of observing a certain mean difference when cell group assignments are random. The p-value was then computed based on how the actual mean difference compared to this null distribution. In the bootstrap analysis for Figure 5, we resampled data points with replacement from the original dataset. The 99% confidence intervals were derived from the 0.5^th^ to the 99.5^th^ percentiles of this resampled distribution. To compute a bootstrap-based p-value, we constructed a distribution based on the differences between the resampled distributions of two groups. The p-value was then determined by the proportion of these differences that were above (or below) zero.

### Anatomy

After completion of the electrophysiological recordings, mice were anaesthetized with pentobarbital and transcardially perfused with 4% PFA (150ml, 8 – 10 ml/min). Subsequently, the heads with implanted tetrode microdrives were post-fixed overnight at 4°C. The next day, the implanted microdrives were removed and the brains were resected followed by post-fixation in 4% PFA for 2 hours. Then brains were either sectioned at 50 micrometers on a vibratome (Leica Microsystems VT1000S) or embedded in gelatine and cryoprotected to be sectioned on a freezing microtome (Epredia HM 450 with a Physitemp BFS-40MPA freezing stage). For embedding, brains were first incubated overnight in 10% sucrose 0.1M Phosphate Buffer solution at 4°C. Then embedded in gelatine (12% gelatine / 10% sucrose), stored in 30% sucrose 0.1M PB overnight at 4°C for cryoprotection, and sectioned the following day utilizing the freezing sliding microtome (Stedehouder et al., 2018). Sections were stained with 4’,6-diamidino-2-phenylindole (DAPI; 0.5 μg/ml, Sigma-Aldrich, cat# D8417) diluted in PB to label cell nuclei, mounted, and cover-slipped with Vectashield mounting medium. Images were acquired using a Zeiss LSM 880 confocal microscope equipped with Plan-Apochromat 10x/0.45, 20x/0.8 objectives. DAPI and an empty channel were imaged using excitation wavelengths of 405, and either 458 or 633 nm.

## Supplementary figures, tables, and legends

**Figure S1.**
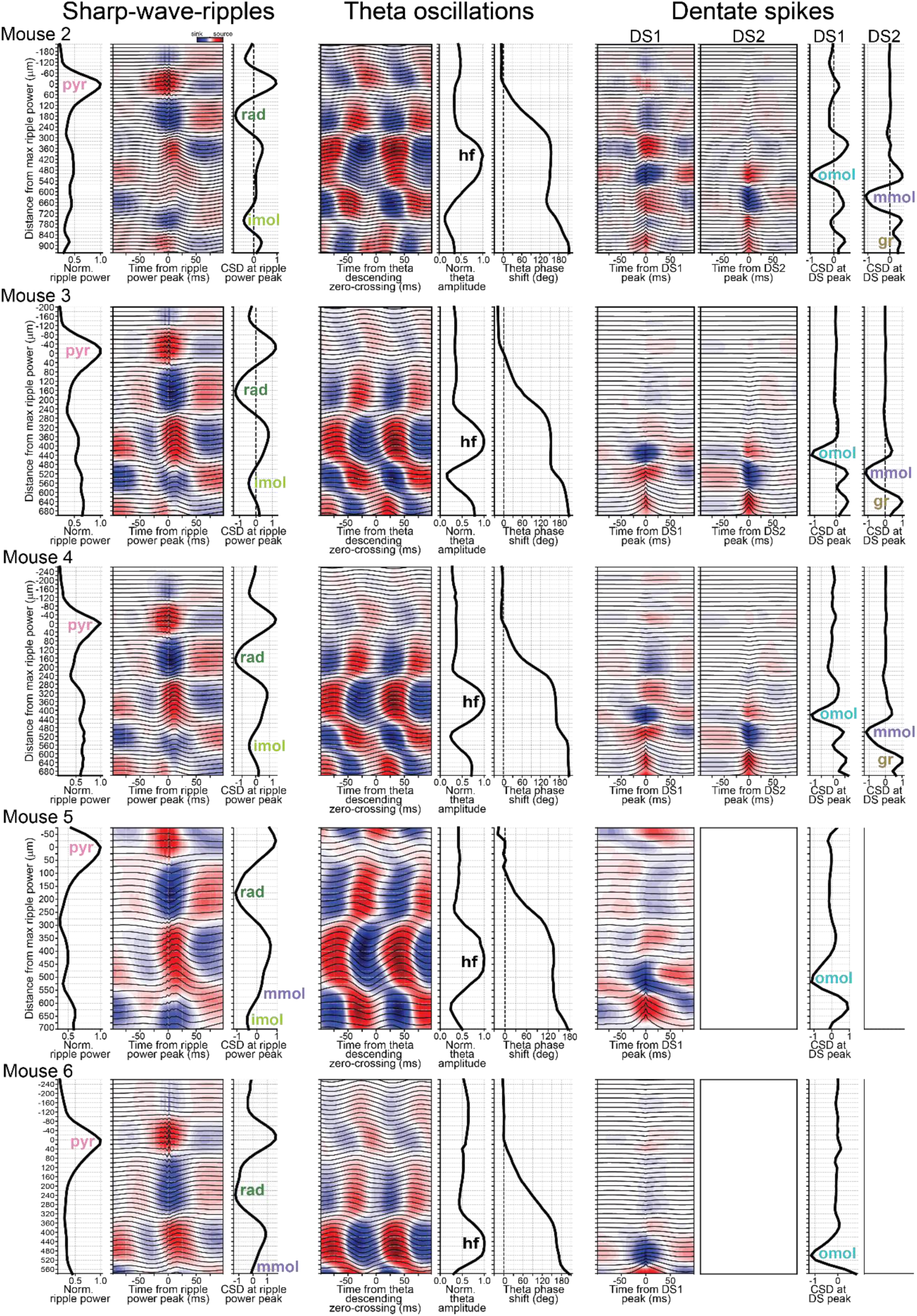
Identification of hippocampal layers using electrophysiological patterns in the silicon probe dataset. Activity profiles used for layer determination in mouse 2 to mouse 6 (see mouse 1 in Figure 1C). From left to right: The sharp-wave ripple panels display ripple power across layers, CSD analysis of LFPs aligned to ripple power peaks, and CSD values at the ripple power peak. The theta oscillation panels display the CSD for LFPs aligned to the pyramidal layer’s descending zero-crossing, alongside normalized theta amplitude and phase shift across layers. The phase shifts are relative to the phase of the pyramidal layer. The dentate spike panels depict CSD analysis for DS1 and DS2 with their respective CSD values at their peak.

**Figure S2.**
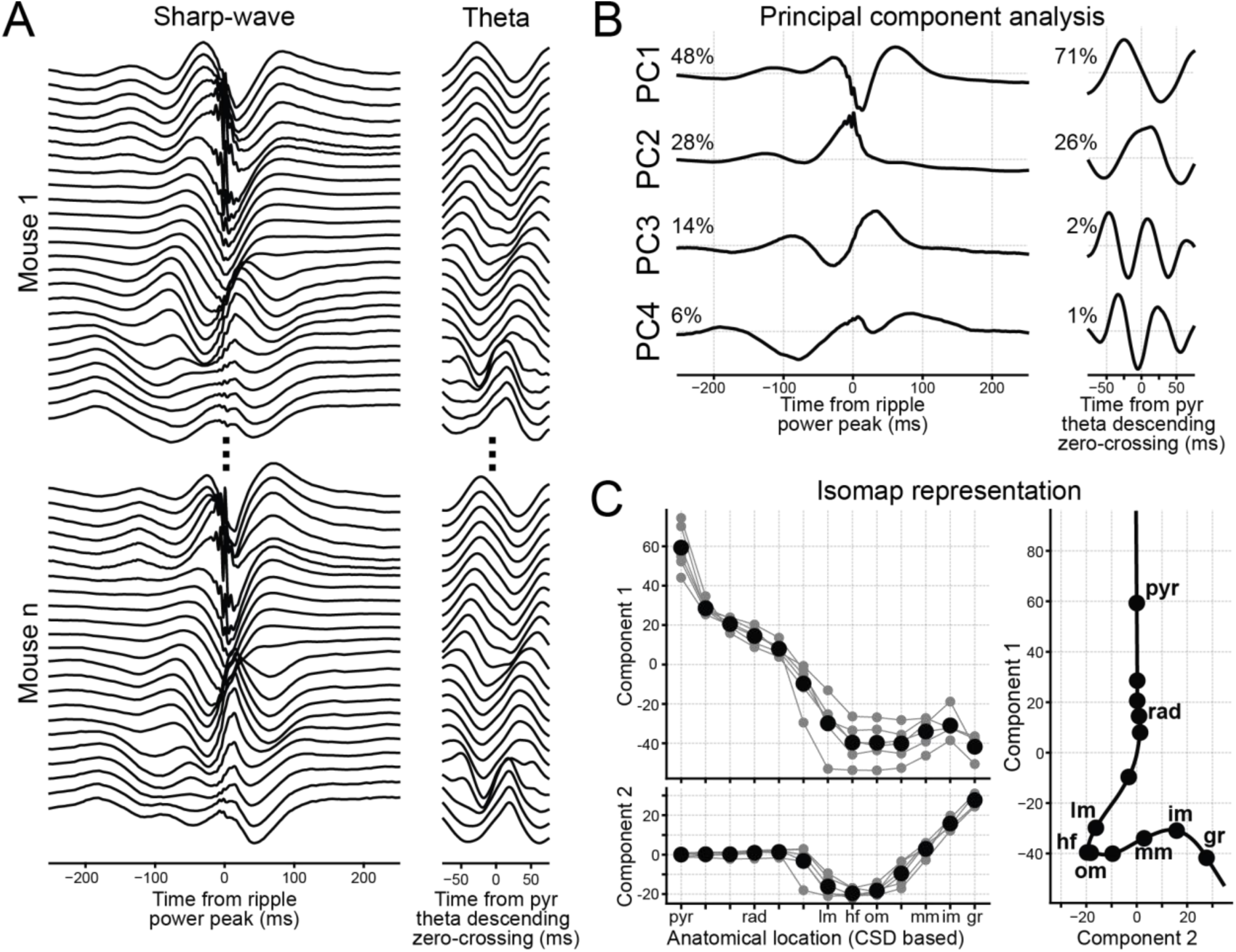
LFP-based feature manifold construction. **(A)** LFP signals employed in defining the feature space. We extracted sharp-wave and theta waveforms from recording channels of the six mice with silicon probe implants. These channels span the dorsal hippocampus radial axis, from the CA1 pyramidal layer to the DG granular layer. **(B)** We then applied principal component analysis to the sharp-wave and theta waveforms. Displayed are the first four principal components (PC 1 to PC 4) for each feature, along with their respective proportion of explained variance. **(C)** Feature traces are then projected onto their first four principal components. Subsequently, a 2-dimensional Isomap is computed based on these feature projections. The left panels depict the feature component coordinates of the CSD-defined layers for each mouse (gray).

**Figure S3.**
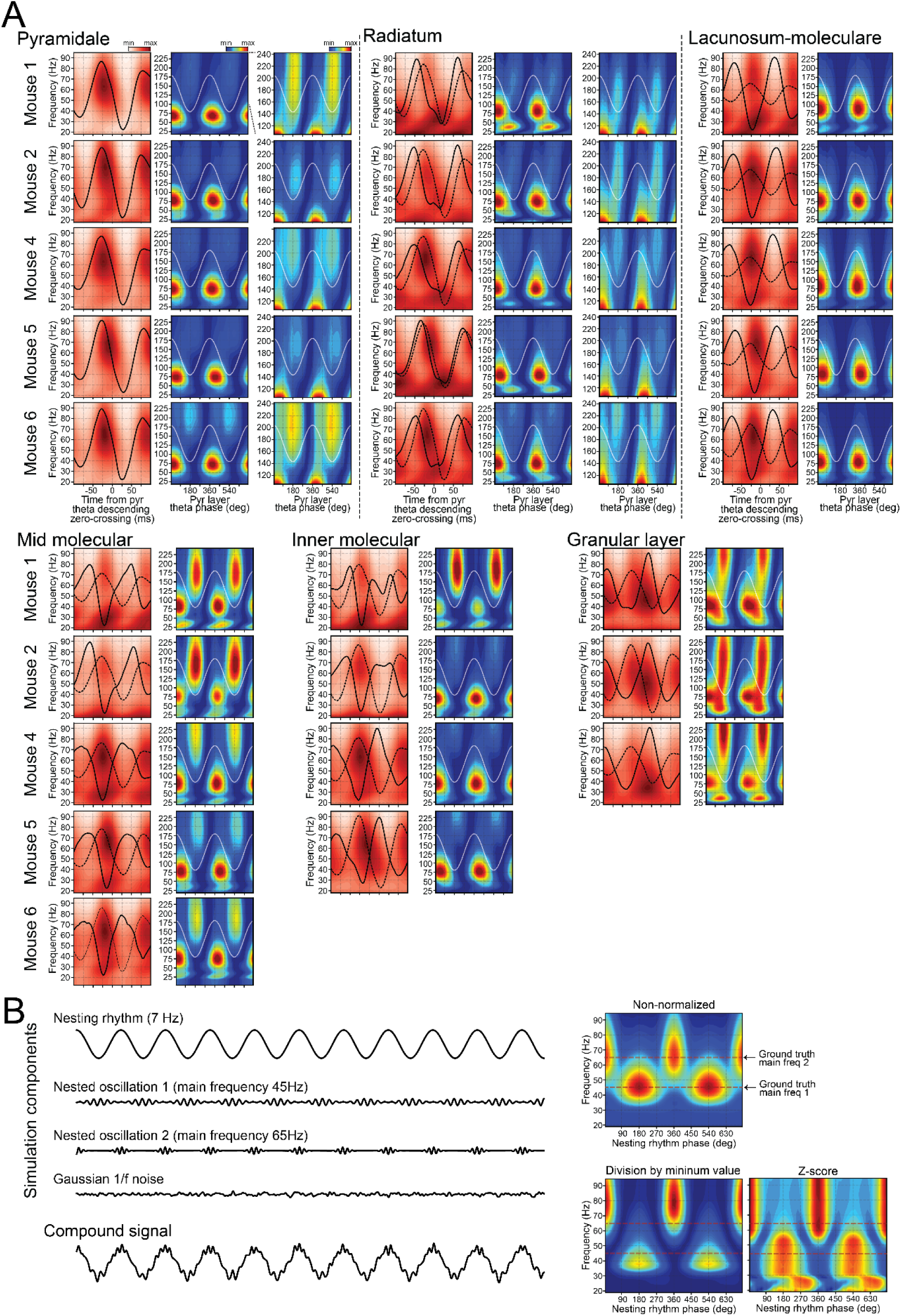
Cross-layer theta-nested gamma profiles for individual mice in the silicon probe dataset. **(A)** Shown are the theta nested-gamma profiles for the other mice in the silicon probe dataset (as in Figure 2). A zoomed-in view into the higher frequencies (> 100 Hz) is provided in the right-most panels for pyramidal and radiatum layers. This magnification distinctly reveals the presence of fast gamma oscillations, which can be partially masked by mid-gamma oscillations. **(B)** Effect of spectrogram normalization on the frequency range for overlapping gamma components. Here, we used *in silicon* data simulation to assess the potential distortion in peak frequency caused by the normalizing frequencies independently. The left traces display the four simulation signal components: a nesting 7-Hz theta-like rhythm, two nested oscillations with known main frequencies, and Gaussian 1/f noise. The nested oscillations are emulated using cosines of defined ground truth frequencies (45 and 65 Hz), with an oscillatory envelope matching the nesting rhythm’s frequency, producing phase-to-amplitude coupling. We then summed these components to generate a simulated compound signal. We finally constructed a theta-gamma-like profile of this compound signal (as in Figure 2). This was achieved without frequency amplitude normalization (top spectrogram), by normalizing each frequency based on its minimum value (bottom left spectrogram), and through z-scoring the amplitude of each frequency (bottom right spectrogram). Notably, the non-normalized version of these approaches offers a more precise portrayal of the foundational frequencies. Dividing frequencies by their minimal value tends to “push apart” the primary frequencies of slightly overlapping components. In contrast, z-scoring amplifies each component’s spectrum tail considerably.

**Figure S4.**
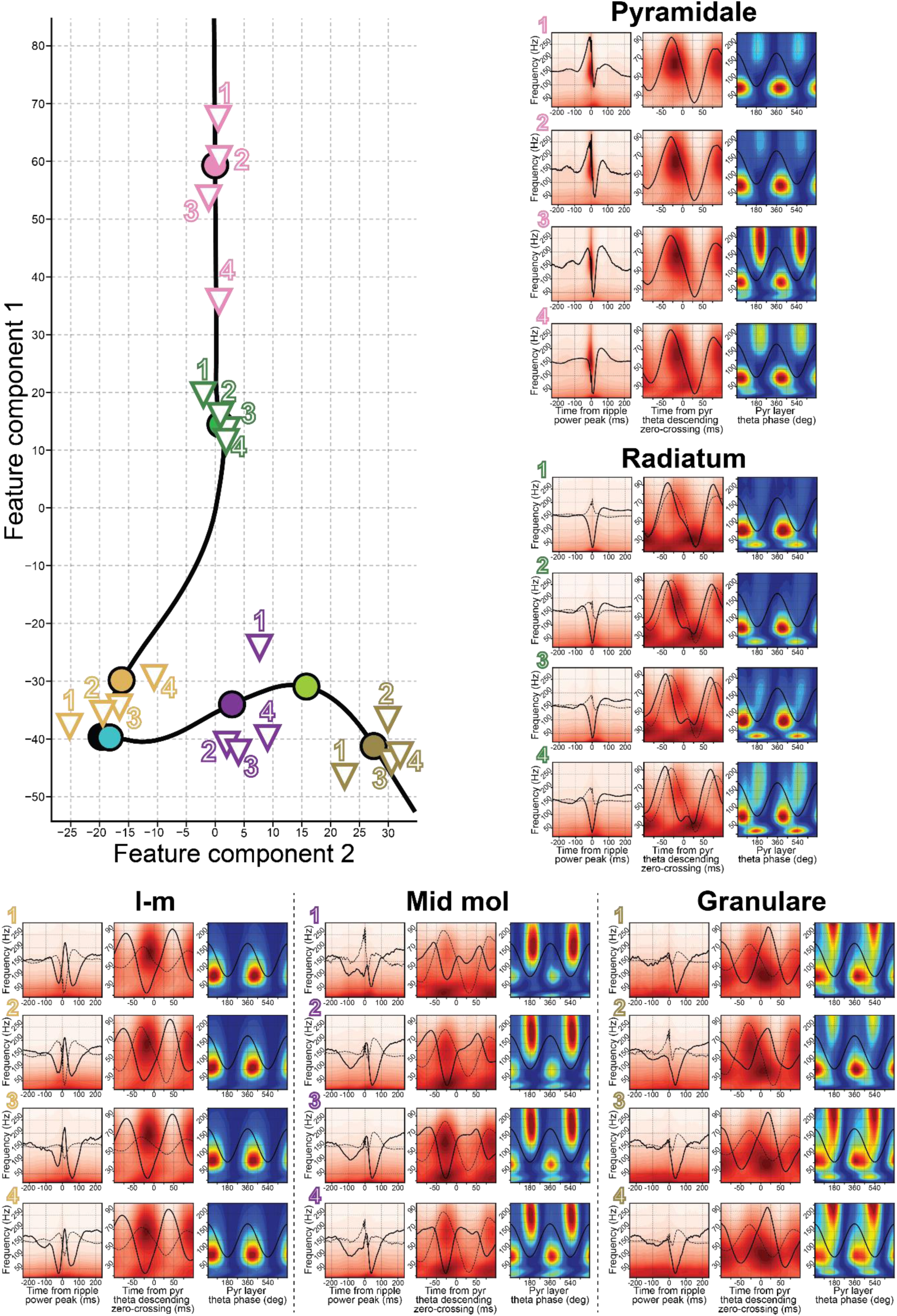
SWR and theta-gamma profiles from tetrode recordings along the CA1-DG axis. This figure features four representative tetrodes, positioned across distinct layers of the CA1-DG axis, guided by the feature manifold. For each tetrode, shown are the SWR and theta-gamma profiles as in Figure 3.

**Figure S5.**
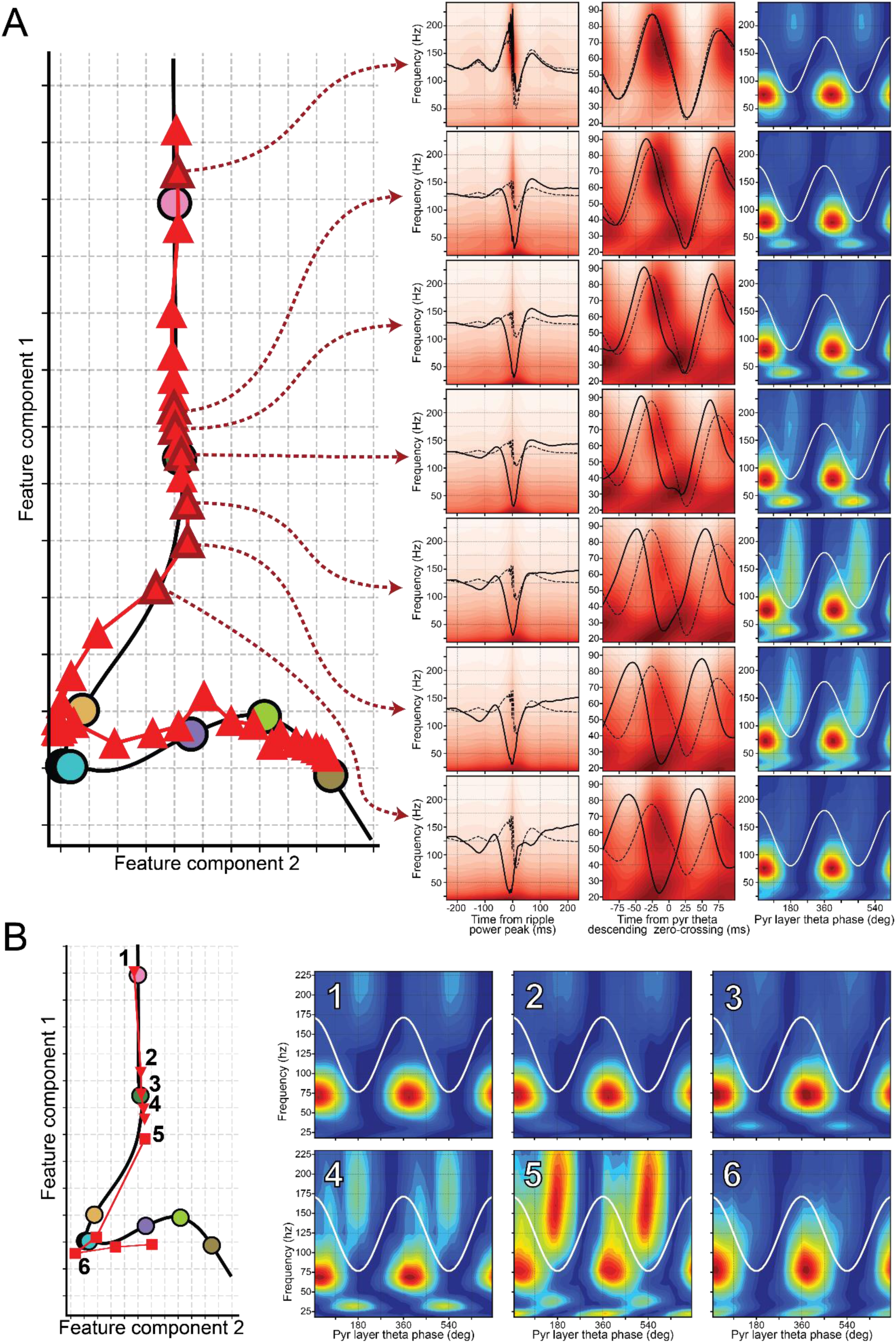
Radiatum fast gamma is observed below the center of the radiatum layer. **(A)** The feature manifold projection of silicon probe channels (spaced at 20 mm) from a representative mouse is displayed. Accompanying this, the SWR and theta-gamma profiles of selected channels are presented on the right, with arrows indicating their respective positions. Notably, the radiatum fast gamma component (visible in the right-most panels showing the normalized theta-gamma profiles) is more pronounced in the recording channel located slightly below the center of the radiatum layer on the manifold. **(B)** A depiction like **A**, but from a tetrode recording experiment as in Figure 3. The experiment began with a tetrode positioned in the pyramidal layer and then gradually moved towards the DG. Theta-gamma profiles for the sessions labeled on the left panel are detailed on the right. Remarkably, during session 5, with the tetrode positioned just below the radiatum center, the radiatum fast gamma component is at its peak clarity.

**Figure S6.**
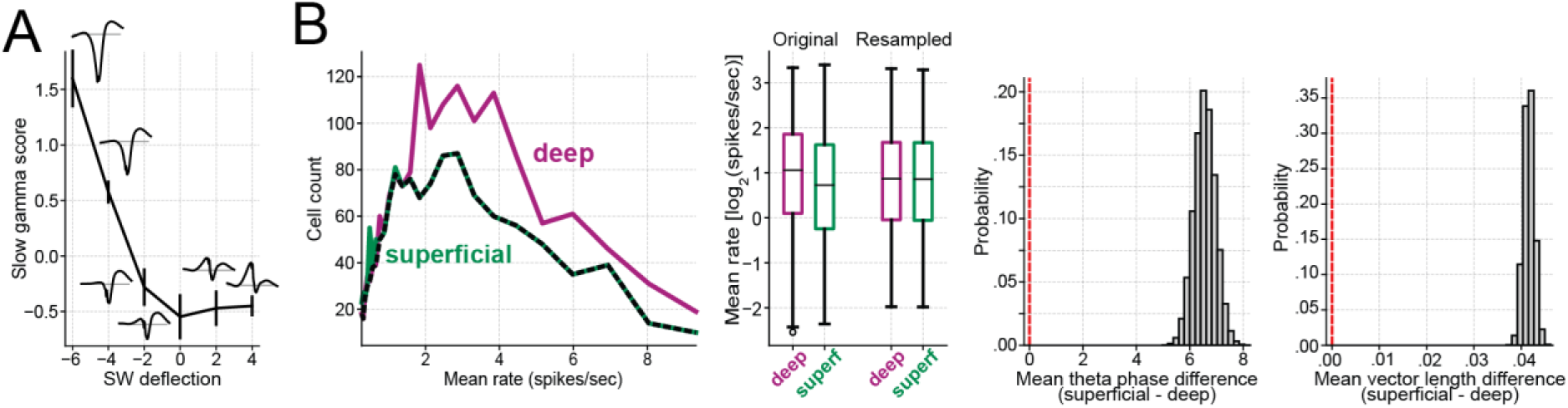
Gamma sharp-wave relationships and rate-matched principal cell analysis. **(A)** Shows the relation between slow gamma amplitude and sharp-wave depth. The slow gamma metric is measured as the amplitude between 25 and 45 Hz during the theta phase of 90 to 270 degrees, divided by its total amplitude above 18 Hz. Sharp-wave deflection is the value obtained from the mean sharp-wave waveform at the ripple power peak, divided by the waveform’s standard deviation. Vertical bars on each data point denote the 99% confidence interval. Each data point is accompanied by insets that represent the average sharp-wave derived from the channels in their corresponding sharp-wave deflection bin. **(B)** Shows the outline of rate-matched analysis for comparing between deep and superficial pyramidal sub-layer cells. The far-left panel shows histograms of average firing rates for each principal cell sub-population, computed over 25 log-spaced bins spanning from 0.25 to 10 spikes per second. The dashed line marks the lowest count across the distributions for each rate bin, guiding the histogram for resampled distributions. The next panel juxtaposes the original rate distributions with a representative resampled version. During resampling, cells are randomly chosen from each rate bin for a designated cell group, aligning with the counts highlighted by the dashed line in the far-left panel, thus assuring that resampled groups have equivalent mean rate distributions. The right-most panels show the distributions of differences in mean theta phase and mean vector length between deep and superficial principal cells across 100,000 resampling operations. These difference distributions consistently stand apart from zero demonstrating disparities in the mean theta phases and the strength of their couplings across the two cell sub-populations even when rate matched.

**Supplementary Table S1.**
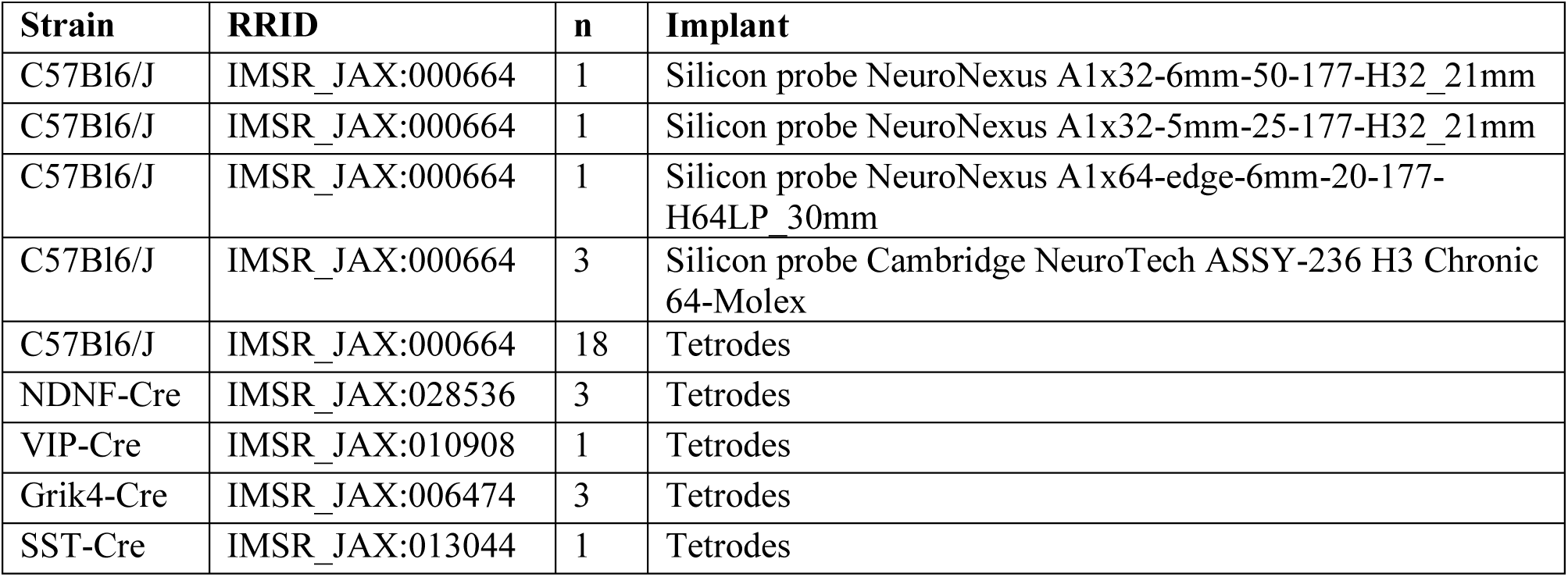
Mice and multichannel recording electrodes.

| Strain | RRID | n | Implant |
| --- | --- | --- | --- |
| C57Bl6/J | IMSR_JAX:000664 | 1 | Silicon probe NeuroNexus A1x32-6mm-50-177-H32_21mm |
| C57Bl6/J | IMSR_JAX:000664 | 1 | Silicon probe NeuroNexus A1x32-5mm-25-177-H32_21mm |
| C57Bl6/J | IMSR_JAX:000664 | 1 | Silicon probe NeuroNexus A1x64-edge-6mm-20-177-H64LP_30mm |
| C57Bl6/J | IMSR_JAX:000664 | 3 | Silicon probe Cambridge NeuroTech ASSY-236 H3 Chronic 64-Molex |
| C57Bl6/J | IMSR_JAX:000664 | 18 | Tetrodes |
| NDNF-Cre | IMSR_JAX:028536 | 3 | Tetrodes |
| VIP-Cre | IMSR_JAX:010908 | 1 | Tetrodes |
| Grik4-Cre | IMSR_JAX:006474 | 3 | Tetrodes |
| SST-Cre | IMSR_JAX:013044 | 1 | Tetrodes |

## References

1. Aery Jones, E.A., Rao, A., Zilberter, M., Djukic, B., Bant, J.S., Gillespie, A.K., Koutsodendris, N., Nelson, M., Yoon, S.Y., Huang, K., Yuan, H., Gill, T.M., Huang, Y., Frank, L.M., 2021. Dentate gyrus and CA3 GABAergic interneurons bidirectionally modulate signatures of internal and external drive to CA1. Cell Reports 37, 110159. 10.1016/j.celrep.2021.110159

2. Andersen, P. (Ed.), 2007. The hippocampus book. Oxford University Press, Oxford ; New York.

3. Belluscio, M.A., Mizuseki, K., Schmidt, R., Kempter, R., Buzsaki, G., 2012. Cross-Frequency Phase-Phase Coupling between Theta and Gamma Oscillations in the Hippocampus. Journal of Neuroscience 32, 423–435. 10.1523/JNEUROSCI.4122-11.2012

4. Benchenane, K., Peyrache, A., Khamassi, M., Tierney, P.L., Gioanni, Y., Battaglia, F.P., Wiener, S.I., 2010. Coherent theta oscillations and reorganization of spike timing in the hippocampal-prefrontal network upon learning. Neuron 66, 921–936. 10.1016/j.neuron.2010.05.013

5. Bragin, A., Jandó, G., Nádasdy, Z., Hetke, J., Wise, K., Buzsáki, G., 1995a. Gamma (40-100 Hz) oscillation in the hippocampus of the behaving rat. J. Neurosci. 15, 47–60.

6. Bragin, A., Jando, G., Nadasdy, Z., van Landeghem, M., Buzsaki, G., 1995b. Dentate EEG spikes and associated interneuronal population bursts in the hippocampal hilar region of the rat. Journal of Neurophysiology 73, 1691–1705. 10.1152/jn.1995.73.4.1691

7. Brankack, J., Stewart, M., Fox, S.E., 1993. Current source density analysis of the hippocampal theta rhythm: associated sustained potentials and candidate synaptic generators. Brain Res. 615, 310– 327.

8. Buzsáki, G., 2015. Hippocampal sharp wave-ripple: A cognitive biomarker for episodic memory and planning. Hippocampus 25, 1073–1188. 10.1002/hipo.22488

9. Buzsáki, G., 2010. Neural Syntax: Cell Assemblies, Synapsembles, and Readers. Neuron 68, 362–385. 10.1016/j.neuron.2010.09.023

10. Buzsáki, G., 2005. Theta rhythm of navigation: Link between path integration and landmark navigation, episodic and semantic memory. Hippocampus 15, 827–840. 10.1002/hipo.20113

11. Buzsáki, G., 1986. Hippocampal sharp waves: their origin and significance. Brain Res. 398, 242–252. 10.1016/0006-8993(86)91483-6

12. Buzsáki, G., Anastassiou, C.A., Koch, C., 2012. The origin of extracellular fields and currents — EEG, ECoG, LFP and spikes. Nat Rev Neurosci 13, 407–420. 10.1038/nrn3241

13. Buzsáki, G., Czopf, J., Kondákor, I., Kellényi, L., 1986. Laminar distribution of hippocampal rhythmic slow activity (RSA) in the behaving rat: Current-source density analysis, effects of urethane and atropine. Brain Research 365, 125–137. 10.1016/0006-8993(86)90729-8

14. Buzsáki, G., Horvath, Z., Urioste, R., Hetke, J., Wise, K., 1992. High-frequency network oscillation in the hippocampus. Science 256, 1025–1027. 10.1126/science.1589772

15. Buzsáki, G., Lai-Wo S., L., Vanderwolf, C.H., 1983. Cellular bases of hippocampal EEG in the behaving rat. Brain Research Reviews 6, 139–171. 10.1016/0165-0173(83)90037-1

16. Buzsáki, G., Wang, X.-J., 2012. Mechanisms of Gamma Oscillations. Annual Review of Neuroscience 35, 203–225. 10.1146/annurev-neuro-062111-150444

17. Colgin, L.L., Denninger, T., Fyhn, M., Hafting, T., Bonnevie, T., Jensen, O., Moser, M.-B., Moser, E.I., 2009. Frequency of gamma oscillations routes flow of information in the hippocampus. Nature 462, 353–357. 10.1038/nature08573

18. Csicsvari, tJozsef, Hirase, H., Czurkó, A., Mamiya, A., Buzsáki, G., 1999. Fast Network Oscillations in the Hippocampal CA1 Region of the Behaving Rat. J. Neurosci. 19, RC20–RC20. 10.1523/JNEUROSCI.19-16-j0001.1999

19. Deering, R., Kaiser, J.F., 2005. The Use of a Masking Signal to Improve Empirical Mode Decomposition, in: Proceedings. (ICASSP ’05). IEEE International Conference on Acoustics, Speech, and Signal Processing, 2005. Presented at the (ICASSP ’05). IEEE International Conference on Acoustics, Speech, and Signal Processing, 2005., IEEE, Philadelphia, Pennsylvania, USA, pp. 485–488. 10.1109/ICASSP.2005.1416051

20. Desmond, N.L., Scott, C.A., Jane, J.A., Levy, W.B., 1994. Ultrastructural identification of entorhinal cortical synapses in CA1 stratum lacunosum-moleculare of the rat. Hippocampus 4, 594–600. 10.1002/hipo.450040509

21. Ego-Stengel, V., Wilson, M.A., 2009. Disruption of ripple-associated hippocampal activity during rest impairs spatial learning in the rat. Hippocampus NA-NA. 10.1002/hipo.20707

22. Eichenbaum, H., 2000. A cortical–hippocampal system for declarative memory. Nat Rev Neurosci 1, 41– 50. 10.1038/35036213

23. El-Gaby, M., Reeve, H.M., Lopes-Dos-Santos, V., Campo-Urriza, N., Perestenko, P.V., Morley, A., Strickland, L.A.M., Lukács, I.P., Paulsen, O., Dupret, D., 2021. An emergent neural coactivity code for dynamic memory. Nat Neurosci 24, 694–704. 10.1038/s41593-021-00820-w

24. Fernández-Ruiz, A., Herreras, O., 2013. Identifying the synaptic origin of ongoing neuronal oscillations through spatial discrimination of electric fields. Front Comput Neurosci 7, 5. 10.3389/fncom.2013.00005

25. Fernández-Ruiz, A., Oliva, A., Nagy, G.A., Maurer, A.P., Berényi, A., Buzsáki, G., 2017. Entorhinal-CA3 Dual-Input Control of Spike Timing in the Hippocampus by Theta-Gamma Coupling. Neuron 93, 1213–1226.e5. 10.1016/j.neuron.2017.02.017

26. Fernández-Ruiz, A., Oliva, A., Soula, M., Rocha-Almeida, F., Nagy, G.A., Martin-Vazquez, G., Buzsáki, G., 2021. Gamma rhythm communication between entorhinal cortex and dentate gyrus neuronal assemblies. Science 372, eabf3119. 10.1126/science.abf3119

27. Fernandez-Ruiz, A., Sirota, A., Lopes-dos-Santos, V., Dupret, D., 2023. Over and above frequency: Gamma oscillations as units of neural circuit operations. Neuron 111, 936–953. 10.1016/j.neuron.2023.02.026

28. Fosso, O.B., Molinas, M., 2018. EMD Mode Mixing Separation of Signals with Close Spectral Proximity in Smart Grids, in: 2018 IEEE PES Innovative Smart Grid Technologies Conference Europe (ISGT-Europe). Presented at the 2018 IEEE PES Innovative Smart Grid Technologies Conference Europe (ISGT-Europe), IEEE, Sarajevo, Bosnia and Herzegovina, pp. 1–6. 10.1109/ISGTEurope.2018.8571816

29. Gava, G.P., McHugh, S.B., Lefèvre, L., Lopes-dos-Santos, V., Trouche, S., El-Gaby, M., Schultz, S.R., Dupret, D., 2021. Integrating new memories into the hippocampal network activity space. Nat Neurosci 24, 326–330. 10.1038/s41593-021-00804-w

30. Gerbrandt, L.K., Fowler, J.R., 1980. Arousal-Related Sustained Potentials in Neocortex and Hippocampus of Rats, in: Progress in Brain Research. Elsevier, pp. 109–116. 10.1016/S0079-6123(08)61614-3

31. Girardeau, G., Benchenane, K., Wiener, S.I., Buzsáki, G., Zugaro, M.B., 2009. Selective suppression of hippocampal ripples impairs spatial memory. Nat Neurosci 12, 1222–1223. 10.1038/nn.2384

32. Girardeau, G., Lopes-dos-Santos, V., 2021. Brain neural patterns and the memory function of sleep. Science 374, 560–564. 10.1126/science.abi8370

33. Green, K.F., Rawlins, J.N.P., 1979. Hippocampal theta in rats under urethane: Generators and phase relations. Electroencephalography and Clinical Neurophysiology 47, 420–429. 10.1016/0013-4694(79)90158-5

34. Grosmark, A.D., Buzsáki, G., 2016. Diversity in neural firing dynamics supports both rigid and learned hippocampal sequences. Science 351, 1440–1443. 10.1126/science.aad1935

35. Hsiao, Y.-T., Zheng, C., Colgin, L.L., 2016. Slow gamma rhythms in CA3 are entrained by slow gamma activity in the dentate gyrus. Journal of Neurophysiology 116, 2594–2603. 10.1152/jn.00499.2016

36. Huang, N.E., Shen, Z., Long, S.R., Wu, M.C., Shih, H.H., Zheng, Q., Yen, N.-C., Tung, C.C., Liu, H.H., 1998. The empirical mode decomposition and the Hilbert spectrum for nonlinear and non-stationary time series analysis. Proceedings of the Royal Society of London. Series A: Mathematical, Physical and Engineering Sciences 454, 903–995. 10.1098/rspa.1998.0193

37. Hyman, J.M., Zilli, E.A., Paley, A.M., Hasselmo, M.E., 2005. Medial prefrontal cortex cells show dynamic modulation with the hippocampal theta rhythm dependent on behavior. Hippocampus 15, 739– 749. 10.1002/hipo.20106

38. Ishizuka, N., Weber, J., Amaral, D.G., 1990. Organization of intrahippocampal projections originating from CA3 pyramidal cells in the rat. J. Comp. Neurol. 295, 580–623. 10.1002/cne.902950407

39. Jones, M.W., Wilson, M.A., 2005. Theta rhythms coordinate hippocampal-prefrontal interactions in a spatial memory task. PLoS Biol. 3, e402. 10.1371/journal.pbio.0030402

40. Karalis, N., Sirota, A., 2022. Breathing coordinates cortico-hippocampal dynamics in mice during offline states. Nat Commun 13, 467. 10.1038/s41467-022-28090-5

41. Kemere, C., Carr, M.F., Karlsson, M.P., Frank, L.M., 2013. Rapid and Continuous Modulation of Hippocampal Network State during Exploration of New Places. PLoS ONE 8, e73114. 10.1371/journal.pone.0073114

42. Király, B., Domonkos, A., Jelitai, M., Lopes-dos-Santos, V., Martínez-Bellver, S., Kocsis, B., Schlingloff, D., Joshi, A., Salib, M., Fiáth, R., Barthó, P., Ulbert, I., Freund, T.F., Viney, T.J., Dupret, D., Varga, V., Hangya, B., 2023. The medial septum controls hippocampal supra-theta oscillations. Nat Commun 14, 6159. 10.1038/s41467-023-41746-0

43. Klausberger, T., Somogyi, P., 2008. Neuronal Diversity and Temporal Dynamics: The Unity of Hippocampal Circuit Operations. Science 321, 53–57. 10.1126/science.1149381

44. Lasztóczi, B., Klausberger, T., 2017. Distinct gamma oscillations in the distal dendritic fields of the dentate gyrus and the CA1 area of mouse hippocampus. Brain Struct Funct 222, 3355–3365. 10.1007/s00429-017-1421-3

45. Lasztóczi, B., Klausberger, T., 2016. Hippocampal Place Cells Couple to Three Different Gamma Oscillations during Place Field Traversal. Neuron 91, 34–40. 10.1016/j.neuron.2016.05.036

46. Lasztóczi, B., Klausberger, T., 2014. Layer-Specific GABAergic Control of Distinct Gamma Oscillations in the CA1 Hippocampus. Neuron 81, 1126–1139. 10.1016/j.neuron.2014.01.021

47. Lee, S.-H., Marchionni, I., Bezaire, M., Varga, C., Danielson, N., Lovett-Barron, M., Losonczy, A., Soltesz, I., 2014. Parvalbumin-Positive Basket Cells Differentiate among Hippocampal Pyramidal Cells. Neuron 82, 1129–1144. 10.1016/j.neuron.2014.03.034

48. Lensu, S., Waselius, T., Penttonen, M., Nokia, M.S., 2019. Dentate spikes and learning: disrupting hippocampal function during memory consolidation can improve pattern separation. Journal of Neurophysiology 121, 131–139. 10.1152/jn.00696.2018

49. Lopes-dos-Santos, V., van de Ven, G.M., Morley, A., Trouche, S., Campo-Urriza, N., Dupret, D., 2018. Parsing Hippocampal Theta Oscillations by Nested Spectral Components during Spatial Exploration and Memory-Guided Behavior. Neuron 100, 940–952.e7. 10.1016/j.neuron.2018.09.031

50. López-Madrona, V.J., Pérez-Montoyo, E., Álvarez-Salvado, E., Moratal, D., Herreras, O., Pereda, E., Mirasso, C.R., Canals, S., 2020. Different theta frameworks coexist in the rat hippocampus and are coordinated during memory-guided and novelty tasks. eLife 9, e57313. 10.7554/eLife.57313

51. Magland, J., Jun, J.J., Lovero, E., Morley, A.J., Hurwitz, C.L., Buccino, A.P., Garcia, S., Barnett, A.H., 2020. SpikeForest, reproducible web-facing ground-truth validation of automated neural spike sorters. eLife 9, e55167. 10.7554/eLife.55167

52. Mitzdorf, U., 1985. Current source-density method and application in cat cerebral cortex: investigation of evoked potentials and EEG phenomena. Physiol. Rev. 65, 37–100. 10.1152/physrev.1985.65.1.37

53. Mizuseki, K., Diba, K., Pastalkova, E., Buzsáki, G., 2011. Hippocampal CA1 pyramidal cells form functionally distinct sublayers. Nat Neurosci 14, 1174–1181. 10.1038/nn.2894

54. Mizuseki, K., Sirota, A., Pastalkova, E., Buzsáki, G., 2009. Theta Oscillations Provide Temporal Windows for Local Circuit Computation in the Entorhinal-Hippocampal Loop. Neuron 64, 267–280. 10.1016/j.neuron.2009.08.037

55. Navas-Olive, A., Valero, M., Jurado-Parras, T., de Salas-Quiroga, A., Averkin, R.G., Gambino, G., Cid, E., de la Prida, L.M., 2020. Multimodal determinants of phase-locked dynamics across deep-superficial hippocampal sublayers during theta oscillations. Nat Commun 11, 2217. 10.1038/s41467-020-15840-6

56. O’Keefe, J., Nadel, L., 1978. The hippocampus as a cognitive map. Clarendon Press ; Oxford University Press, Oxford : New York.

57. Pachitariu, M., Steinmetz, N.A., Kadir, S.N., Carandini, M., Harris, K.D., 2016. Fast and accurate spike sorting of high-channel count probes with KiloSort, in: Lee, D., Sugiyama, M., Luxburg, U., Guyon, I., Garnett, R. (Eds.), Advances in Neural Information Processing Systems. Curran Associates, Inc.

58. Penttonen, M., Kamondi, A., Sik, A., Acsády, L., Buzsáki, G., 1997. Feed-forward and feed-back activation of the dentate gyrus in vivo during dentate spikes and sharp wave bursts. Hippocampus 7, 437– 450. 10.1002/(SICI)1098-1063(1997)7:4<437::AID-HIPO9>3.0.CO;2-F

59. Quinn, A., Lopes-dos-Santos, V., Dupret, D., Nobre, A., Woolrich, M., 2021a. EMD: Empirical Mode Decomposition and Hilbert-Huang Spectral Analyses in Python. JOSS 6, 2977. 10.21105/joss.02977

60. Quinn, A., Lopes-dos-Santos, V., Huang, N., Liang, W.-K., Juan, C.-H., Yeh, J.-R., Nobre, A.C., Dupret, D., Woolrich, M.W., 2021b. Within-cycle instantaneous frequency profiles report oscillatory waveform dynamics. Journal of Neurophysiology 126, 1190–1208. 10.1152/jn.00201.2021

61. Sakalar, E., Klausberger, T., Lasztóczi, B., 2022. Neurogliaform cells dynamically decouple neuronal synchrony between brain areas. Science 377, 324–328. 10.1126/science.abo3355

62. Scheffer-Teixeira, R., Belchior, H., Caixeta, F.V., Souza, B.C., Ribeiro, S., Tort, A.B.L., 2012. Theta phase modulates multiple layer-specific oscillations in the CA1 region. Cereb. Cortex 22, 2404–2414. 10.1093/cercor/bhr319

63. Scheffer-Teixeira, R., Belchior, H., Leão, R.N., Ribeiro, S., Tort, A.B.L., 2013. On high-frequency field oscillations (>100 Hz) and the spectral leakage of spiking activity. J. Neurosci. 33, 1535–1539. 10.1523/JNEUROSCI.4217-12.2013

64. Schomburg, E.W., Fernández-Ruiz, A., Mizuseki, K., Berényi, A., Anastassiou, C.A., Koch, C., Buzsáki, G., 2014. Theta phase segregation of input-specific gamma patterns in entorhinal-hippocampal networks. Neuron 84, 470–485. 10.1016/j.neuron.2014.08.051

65. Senzai, Y., Buzsáki, G., 2017. Physiological Properties and Behavioral Correlates of Hippocampal Granule Cells and Mossy Cells. Neuron 93, 691–704.e5. 10.1016/j.neuron.2016.12.011

66. Siapas, A.G., Lubenov, E.V., Wilson, M.A., 2005. Prefrontal Phase Locking to Hippocampal Theta Oscillations. Neuron 46, 141–151. 10.1016/j.neuron.2005.02.028

67. Sirota, A., Buzsáki, G., 2005. Interaction between neocortical and hippocampal networks via slow oscillations. THL 3, 245. 10.1017/S1472928807000258

68. Sirota, A., Csicsvari, J., Buhl, D., Buzsaki, G., 2003. Communication between neocortex and hippocampus during sleep in rodents. Proceedings of the National Academy of Sciences 100, 2065–2069. 10.1073/pnas.0437938100

69. Sirota, A., Montgomery, S., Fujisawa, S., Isomura, Y., Zugaro, M., Buzsáki, G., 2008. Entrainment of neocortical neurons and gamma oscillations by the hippocampal theta rhythm. Neuron 60, 683– 697. 10.1016/j.neuron.2008.09.014

70. Soltesz, I., Losonczy, A., 2018. CA1 pyramidal cell diversity enabling parallel information processing in the hippocampus. Nat Neurosci 21, 484–493. 10.1038/s41593-018-0118-0

71. Someck, S., Levi, A., Sloin, H.E., Spivak, L., Gattegno, R., Stark, E., 2023. Positive and biphasic extracellular waveforms correspond to return currents and axonal spikes. Commun Biol 6, 950. 10.1038/s42003-023-05328-6

72. Spruston, N., 2008. Pyramidal neurons: dendritic structure and synaptic integration. Nat Rev Neurosci 9, 206–221. 10.1038/nrn2286

73. Stark, E., Roux, L., Eichler, R., Senzai, Y., Royer, S., Buzsáki, G., 2014. Pyramidal Cell-Interneuron Interactions Underlie Hippocampal Ripple Oscillations. Neuron 83, 467–480. 10.1016/j.neuron.2014.06.023

74. Stedehouder, J., Brizee, D., Shpak, G., Kushner, S.A., 2018. Activity-Dependent Myelination of Parvalbumin Interneurons Mediated by Axonal Morphological Plasticity. J. Neurosci. 38, 3631– 3642. 10.1523/JNEUROSCI.0074-18.2018

75. Steriade, M., Nunez, A., Amzica, F., 1993. A novel slow (< 1 Hz) oscillation of neocortical neurons in vivo: depolarizing and hyperpolarizing components. J. Neurosci. 13, 3252–3265. 10.1523/JNEUROSCI.13-08-03252.1993

76. Sullivan, D., Csicsvari, J., Mizuseki, K., Montgomery, S., Diba, K., Buzsaki, G., 2011. Relationships between Hippocampal Sharp Waves, Ripples, and Fast Gamma Oscillation: Influence of Dentate and Entorhinal Cortical Activity. Journal of Neuroscience 31, 8605–8616. 10.1523/JNEUROSCI.0294-11.2011

77. Tort, A.B.L., Komorowski, R.W., Manns, J.R., Kopell, N.J., Eichenbaum, H., 2009. Theta-gamma coupling increases during the learning of item-context associations. Proceedings of the National Academy of Sciences 106, 20942–20947. 10.1073/pnas.0911331106

78. Valero, M., Averkin, R.G., Fernandez-Lamo, I., Aguilar, J., Lopez-Pigozzi, D., Brotons-Mas, J.R., Cid, E., Tamas, G., Menendez de la Prida, L., 2017. Mechanisms for Selective Single-Cell Reactivation during Offline Sharp-Wave Ripples and Their Distortion by Fast Ripples. Neuron 94, 1234–1247.e7. 10.1016/j.neuron.2017.05.032

79. Valero, M., Cid, E., Averkin, R.G., Aguilar, J., Sanchez-Aguilera, A., Viney, T.J., Gomez-Dominguez, D., Bellistri, E., de la Prida, L.M., 2015. Determinants of different deep and superficial CA1 pyramidal cell dynamics during sharp-wave ripples. Nat Neurosci 18, 1281–1290. 10.1038/nn.4074

80. Valero, M., de la Prida, L.M., 2018. The hippocampus in depth: a sublayer-specific perspective of entorhinal–hippocampal function. Current Opinion in Neurobiology, Systems Neuroscience 52, 107–114. 10.1016/j.conb.2018.04.013

81. van de Ven, G.M., Trouche, S., McNamara, C.G., Allen, K., Dupret, D., 2016. Hippocampal Offline Reactivation Consolidates Recently Formed Cell Assembly Patterns during Sharp Wave-Ripples. Neuron 92, 968–974. 10.1016/j.neuron.2016.10.020

82. Van Groen, T., Miettinen, P., Kadish, I., 2003. The entorhinal cortex of the mouse: Organization of the projection to the hippocampal formation. Hippocampus 13, 133–149. 10.1002/hipo.10037

83. Van Strien, N.M., Cappaert, N.L.M., Witter, M.P., 2009. The anatomy of memory: an interactive overview of the parahippocampal–hippocampal network. Nat Rev Neurosci 10, 272–282. 10.1038/nrn2614

84. Vanderwolf, C.H., 1969. Hippocampal electrical activity and voluntary movement in the rat. Electroencephalography and Clinical Neurophysiology 26, 407–418. 10.1016/0013-4694(69)90092-3

85. Wang, X.-J., 2010. Neurophysiological and Computational Principles of Cortical Rhythms in Cognition. Physiological Reviews 90, 1195–1268. 10.1152/physrev.00035.2008

86. Whittington, M.A., Traub, R.D., Kopell, N., Ermentrout, B., Buhl, E.H., 2000. Inhibition-based rhythms: experimental and mathematical observations on network dynamics. Int J Psychophysiol 38, 315– 336.

87. Witter, M.P., Wouterlood, F.G., Naber, P.A., Van Haeften, T., 2000. Anatomical organization of the parahippocampal-hippocampal network. Ann. N. Y. Acad. Sci. 911, 1–24.

88. Ylinen, A., Bragin, A., Nádasdy, Z., Jandó, G., Szabó, I., Sik, A., Buzsáki, G., 1995. Sharp wave-associated high-frequency oscillation (200 Hz) in the intact hippocampus: network and intracellular mechanisms. J. Neurosci. 15, 30–46.

